# The *Chlamydia* effector IncE employs two short linear motifs to reprogram host vesicle trafficking

**DOI:** 10.1101/2024.04.23.590830

**Authors:** Khavong Pha, Katherine Mirrashidi, Jessica Sherry, Cuong Joseph Tran, Clara M. Herrera, Eleanor McMahon, Cherilyn A. Elwell, Joanne N. Engel

## Abstract

*Chlamydia trachomatis,* a leading cause of bacteria sexually transmitted infections, creates a specialized intracellular replicative niche by translocation and insertion of a diverse array of effectors (Incs) into the inclusion membrane. Here, we characterize IncE, a multi-functional Inc that encodes two non-overlapping short linear motifs (SLiMs) within its short cytosolic C-terminus. The proximal SLiM mimics an R-SNARE motif to recruit syntaxin (STX) 7 and 12-containing vesicles to the inclusion. The distal SLiM mimics the Sorting Nexin (SNX) 5 and 6 cargo binding site to recruit SNX6-containing vesicles to the inclusion. By simultaneously binding to two distinct vesicle classes, IncE reprograms host cell trafficking to promote the formation of a class of hybrid vesicles at the inclusion that are required for *C. trachomatis* intracellular development. Our work suggests that Incs may have evolved SLiMs to facilitate rapid evolution in a limited protein space to disrupt host cell processes.

## Introduction

*Chlamydia trachomatis (Ct)* infections are important causes of human disease for which no vaccine exists. Although infections can be treated with antibiotics, no drug is cost-effective enough for widespread elimination of disease in developing nations (Hocking et al., 2023). This obligate intracellular parasite replicates within a privileged niche--a membrane bound compartment termed the inclusion—where it subverts multiple host cell functions to survive (Elwell et al., 2016). *Chlamydiae* devotes up to 10% of its genome to a unique family of secreted effectors, the Incs (<u>Inc</u>lusion membrane proteins), which are translocated from the bacteria through the type III secretion system and inserted into the inclusion membrane (Rockey et al., 2002). These effectors are ideally positioned at the host-pathogen interface to function as scaffolds to mediate interactions between the inclusion and the host. We previously employed high throughput affinity purification-mass spectroscopy (AP-MS) of human cells transfected with Incs to identify their host binding partners, with the goal of uncovering the function of Incs (Mirrashidi et al., 2015). This analysis identified predicted high confidence interactions between IncE, an early expressed Inc of unknown function, and proteins associated with two unrelated vesicle classes: (i) sorting nexins (SNX)5 and SNX6 and (ii) syntaxins (STX)7 and STX12. SNX5 and SNX6, together with SNX1 and SNX2, comprise a subclass of retromer, the ESCPE-1 complex (Simonetti et al., 2019). This multiprotein complex mediates retrograde-dependent recycling of cargo receptors from the endosome to the trans golgi network (TGN) (Seaman, 2021; Simonetti et al., 2019). We and others solved the crystal structure of the IncE:SNX5 complex (Elwell et al., 2017; Paul et al., 2017; Sun et al., 2017; Yong et al., 2020). We discovered that the C-terminal 24 amino acids of IncE bind with high affinity directly to a phylogenetically conserved hydrophobic groove in SNX5, which is also conserved in SNX6 homologs and is the site at which cargo binds to SNX5 and SNX6 (Simonetti et al., 2019). Simultaneous depletion of SNX5 and SNX6 results in increased production of infectious progeny (Aeberhard et al., 2015; Mirrashidi et al., 2015), suggesting that the ESCPE-1 complex, or at least SNX5 and SNX6, function as restriction factors to limit *Ct* intracellular infection.

In this study, we explore the role of the predicted second class of IncE interactors, syntaxin (STX) 7 and STX12. STX7 and STX12 are closely related Q-SNARE proteins that mediate late and early endosome vesicle fusion, respectively (Collins et al., 2002; West et al., 2021), but their role in *Ct* pathogenesis is unknown. Notably, neither STX7 nor STX12 have previously been associated with either SNX5/6 or the ESCPE-1 complex. STXs encode a 60-70 amino acid SNARE domain that can hetero-multimerize with other SNARE domains (Bonifacino and Glick, 2004; Hong, 2005). Typically, 1 R-SNARE associates with 3 Q-SNARES, distributed between a donor membrane and a target membrane, along with Rab GTPases and the membrane tethering complex. Upon docking, the 4 SNARE domains align and form a stable 4 helix bundle (the trans-SNARE complex) to allow the “zippering” of the inner and outer leaflets of the donor and target membranes, leading to membrane fusion. A key feature of the SNARE domain is a highly conserved central ionic “0” layer that contains either a conserved arginine (R-SNARE) or glutamine (Q-SNARE). The salt bridges formed by the ionic layer, together with multiple hydrophobic interactions contributed by the flanking 7-8 leucine (heptad) repeats, nucleate the zippering that leads to membrane fusion (Fasshauer et al., 1998; Sutton et al., 1998). Our studies reveal that IncE encodes a short linear motif (SLiM) that mimics the −1 and 0 (ionic) layer of some R-SNARES. We demonstrate that these 6 amino acids are required for IncE binding to STX7 and to STX12, supporting our hypothesis that IncE binds to STX7 and to STX12 through molecular mimicry. We found that STX7 and STX12 contribute to distinct steps in the *Ct* intracellular lifecycle. IncE binding to STX7 is required for production of infectious progeny whereas IncE binding to STX12 facilitates homotypic inclusion fusion. We show that IncE binds to SNX5 or SNX6 and either STX7 or STX12 simultaneously. By tethering 2 distinct vesicle classes at or near the inclusion, IncE facilitates the formation of hybrid vesicles that may contribute to the intracellular survival of this medically important pathogen.

## Results

### IncE binds specifically to STX7 and STX12

Our interactome data set identified STX7 and STX12 as high confidence interactors with IncE (Mirrashidi et al., 2015). We confirmed by transfection studies in HEK293T cells that endogenous STX7 and STX12, but not STX2 (which did not co-affinity purify (co-AP) with transfected IncE (Mirrashidi et al., 2015)), co-AP’d with transfected IncE_Strep_ but not with IncG_Strep_, an unrelated Inc (Fig. S1A). In HeLa cells infected with *Ct* strain LGV/L2 (L2) expressing IncE_FLAG_ or IncG_FLAG_ from a plasmid (L2+pIncE or L2+pIncG, respectively), STX7 and STX12, but not STX2, co-AP’d with IncE but not with IncG (Fig. S1B). Together, these results confirm a highly specific interaction between IncE and STX7 and STX12 in two different cell lines.

### IncE binds STX7 and STX12 by mimicking a short motif of the −1 and 0 (ionic) layer of an R-SNARE domain

STX7 and STX12 are Q-SNARES that are more closely related to each other than to other Q SNARES (Fig. S2), suggesting that IncE might bind to a region common to STX7 and STX12. To investigate the regions of STX7 and STX12 that are necessary for binding to IncE and to delineate the region of IncE required for binding to STX7 and STX12, we constructed deletion mutants of STX7, STX12, and IncE (Fig. 1) and tested their ability to co-AP upon co-transfection. This analysis revealed that the SNARE and transmembrane regions of STX7 or of STX12 are required for binding to IncE (Figs. 1A-D). For IncE, amino acids 37-100 are necessary and sufficient for the interaction with endogenous STX7 and STX12 (Figs. 1E,F). We note that the STX7/12 binding region of IncE is distinct from its previously defined SNX5/6 binding region (Fig. S3A) (Elwell et al., 2017; Paul et al., 2017; Sun et al., 2017; Yong et al., 2020).

**Fig. 1.**
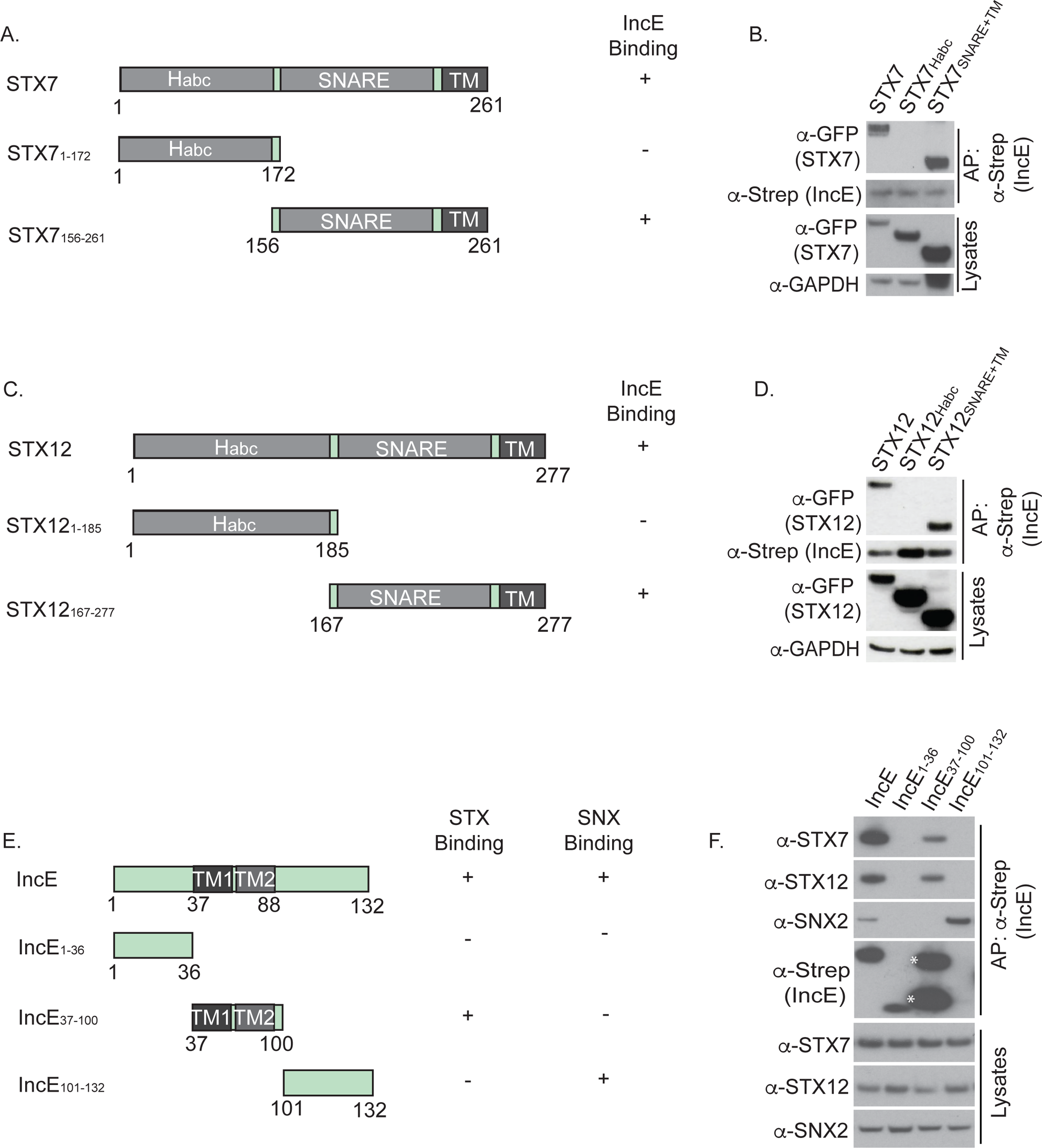
Defining the binding interface of STX7, STX12, and IncE. (A, C, E) Schematic of (A) STX7 (C) STX12, and (E) IncE constructs. H_abc_, regulatory domain; TM, transmembrane. (B, D) Co-AP of IncE_Strep_ with (B) GFP-STX7 or (D) GFP-STX12 deletion constructs. GAPDH serves as a loading control. (F) Co-affinity purification of IncE_Strep_ constructs with STX7, STX12, or SNX2. IncE monomers and dimers are indicated by asterisk. SNX2 serves as a loading control as well as a positive control for SNX binding to IncE. Data shown are representative of two biological replicates.

We further refined the region of IncE required for STX7/12 binding. As residues 37-100 encompasses both transmembrane (TM) domains (Fig. S3B), we considered that the residues C-terminal to the second TM domain might encode the key STX7/12 interacting residues. Indeed, we identified a 6-amino acid sequence (VLENHG) that is homologous to the −1 and 0 (ionic) layer regions of two R-SNARES, human VAMP3 (VLERDQ) and mouse VAMP5 (VLERHG). These 6-amino acid sequence is conserved, albeit with some variation, in IncE homologs (Fig. S3C), suggesting that it is important to IncE function. We hypothesized that this region of IncE may mimic an R-SNARE motif to bind to the STX7 and STX12 Q-SNAREs.

To directly address the role of the IncE VLENHG sequence (hereafter referred to as the STX binding motif (StxBM)), we changed these 6 residues to alanine (IncE_ΔStxBM_) (Fig. S3D) and tested the ability of this IncE variant to co-AP endogenous STX7 and STX12 in transfection and infection studies. We also created an IncE mutant in which residues V114 and F116 are changed to asparagine and aspartic acid, respectively (Fig. S3D), as these residues are predicted critical binding residues to SNX5 and SNX6, based on our published crystal structure (Elwell et al., 2017). This variant is termed IncE_ΔSnxBM_.

In transfection experiments, IncE_ΔStxBM_ no longer co-AP’d STX7 and STX12 but retained the ability to co-AP SNX5 and SNX6 (Fig. S3E). Conversely, IncE_ΔSnxBM_ failed to co-AP SNX5 and SX6 but still co-AP’d STX7 and STX12 (Fig. S3E). We conclude that both IncE variants fold properly and that the StxBM and SnxBM are required for binding to STX7/12 and SNX5/6, respectively.

To interrogate the role of the IncE StxBM and SnxBM, we generated a non-polar *incE* mutant strain (L2Δ*incE*) and complemented it with the following plasmids encoding anhydrous tetracycline (aTC) inducible FLAG-tagged IncE variants: WT IncE, IncE_ΔStxBM_, or IncE_ΔSnxBM_ (Fig. S3D, also see Fig. S4 for strain construction and characterization of L2Δ*incE*). We confirmed by immunofluorescence microscopy that all IncE variants localized to the inclusion membrane (Fig S5). Consistent with the transfection studies (Fig. S2E), in cells infected with the Ct IncE mutant complmented with the IncE variant that cannot bind STX7 and STX12 (L2Δ*incE*+pIncE_ΔStxBM_), STX7 and STX12 did not co-AP with IncE_ΔStxBM._ Importantly, SNX5 and SNX6 still co-AP’d with IncE_ΔStxBM_ (Figs. 2A,B). Conversely, in cells infected with L2Δ*incE* expressing the IncE variant which cannot bind SNX5 and SNX6 (L2Δ*incE*+pIncE_ΔSnxBM_), SNX5 and SNX6 failed to co-AP with IncE_ΔSnxBM_ whereas STX7 and STX12 still co-AP’d with this IncE variant (Figs. 2A,B). As expected, WT IncE co-AP’d with STX7, STX12, SNX5, and SNX6 (Figs. 2A,B). We conclude that the IncE StxBM is required for binding to STX7 and STX12 but not to SNX5 or SNX6. Conversely, the IncE SnxBM is required for binding to SNX5 and SNX6 but not to STX7 or STX12. These results further underscore that the StxBM is distinct and separate from the IncE SnxBM (Fig. S3A).

**Fig. 2.**
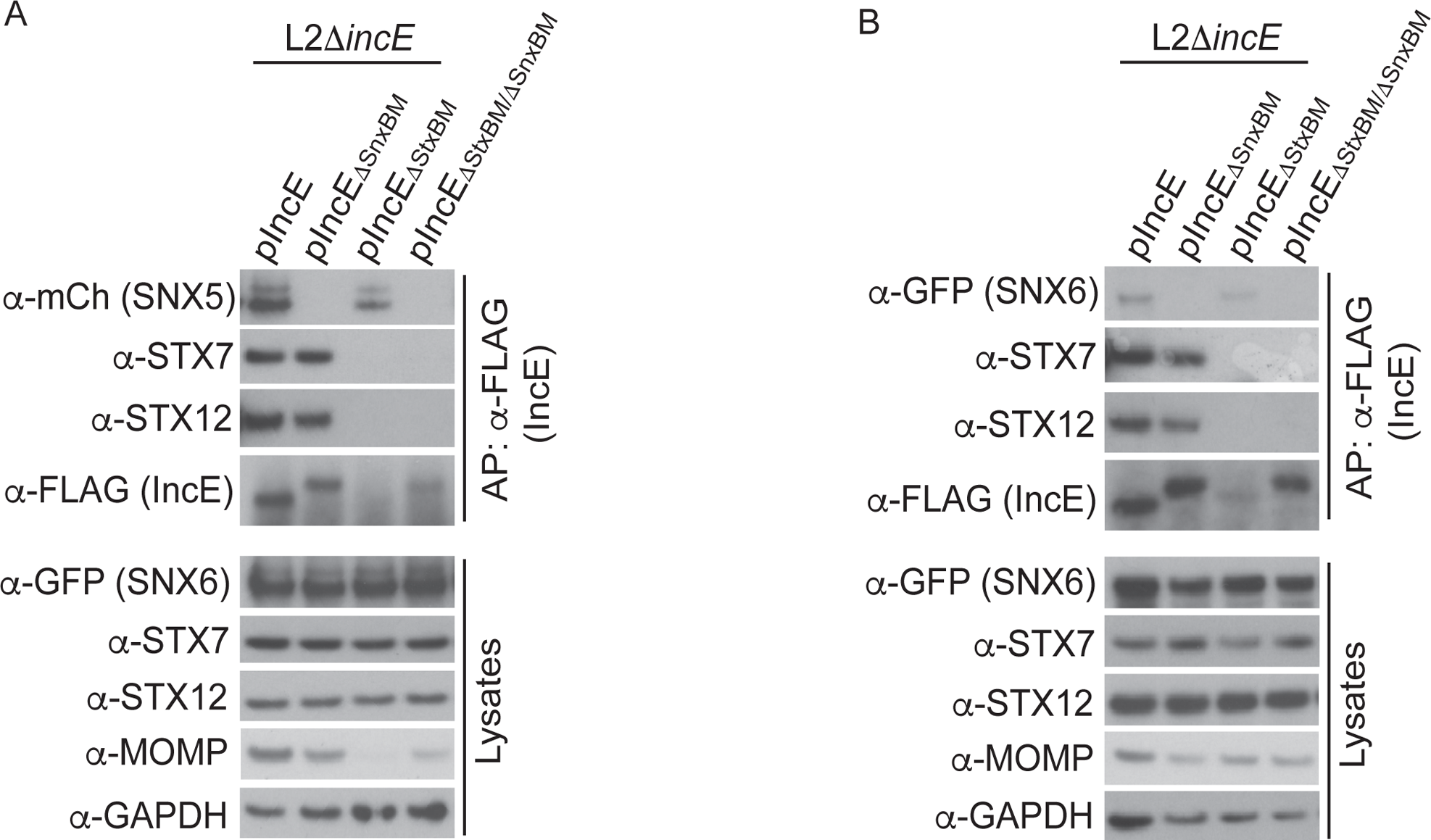
The IncE StxBM and SnxBM are non-overlapping. Co-APs of the indicated IncE constructs with (A) mCherry-SNX5, STX7, and STX12 or with (B) GFP-SNX6, STX7, and STX12. Data shown are representative of immunoblots from three biological replicates.

### IncE binding to STX7 and STX12 is required for late steps of infection

We tested the role of IncE binding to STX7 and STX12 in the pathogenesis of *Ct* infections. Quantitation of *Ct* inclusion formation, specifically the number, size, and localization of inclusions, is a robust metric for early steps in the infection process. Quantification of progeny production additionally assesses late steps in infection (Fig. S6A).

We first quantified inclusion formation and production of infectious progeny by inter-strain comparison of WT (L2+vector) to the *incE* mutant (L2Δ*incE*+vector). Surprisingly, no differences were observed between WT and the *incE* mutant in inclusion formation, inclusion size, or production of infectious progeny (Figs. S6B-D). As depletion of SNX5 and SNX6 enhances progeny production, (Aeberhard et al., 2015; Mirrashidi et al., 2015), if STX7 and/or STX12 are required for progeny production, then the *incE* deletion mutant might exhibit no change in production of infectious progeny compared to WT. With this rationale in mind, we compared progeny production when cells were infected with the individual complemented L2Δ*incE* strains. As the variants are under control of an aTC-inducible promoter, we could perform intra-strain comparisons in the presence or absence of inducer. This strategy has the important advantage that it bypasses the requirement that infection of different strains be performed at equal MOIs, although similar MOIs were used for each strain. Remarkably, cells infected with L2Δ*incE*+pIncE_ΔStxBM_, the IncE variant that cannot bind STX7/12 but that can still bind SNX5/6, exhibited decreased production of infectious progeny (Figs. 3A,B). As well, there was a trend towards decreased inclusion formation in this strain, although it did not reach statistical significance (Figs. 3A,B). Together, these studies reveal a critical role for IncE binding to STXs during intracellular infection that is only unmasked when IncE is still able to bind to SNX5/6.

**Fig. 3.**
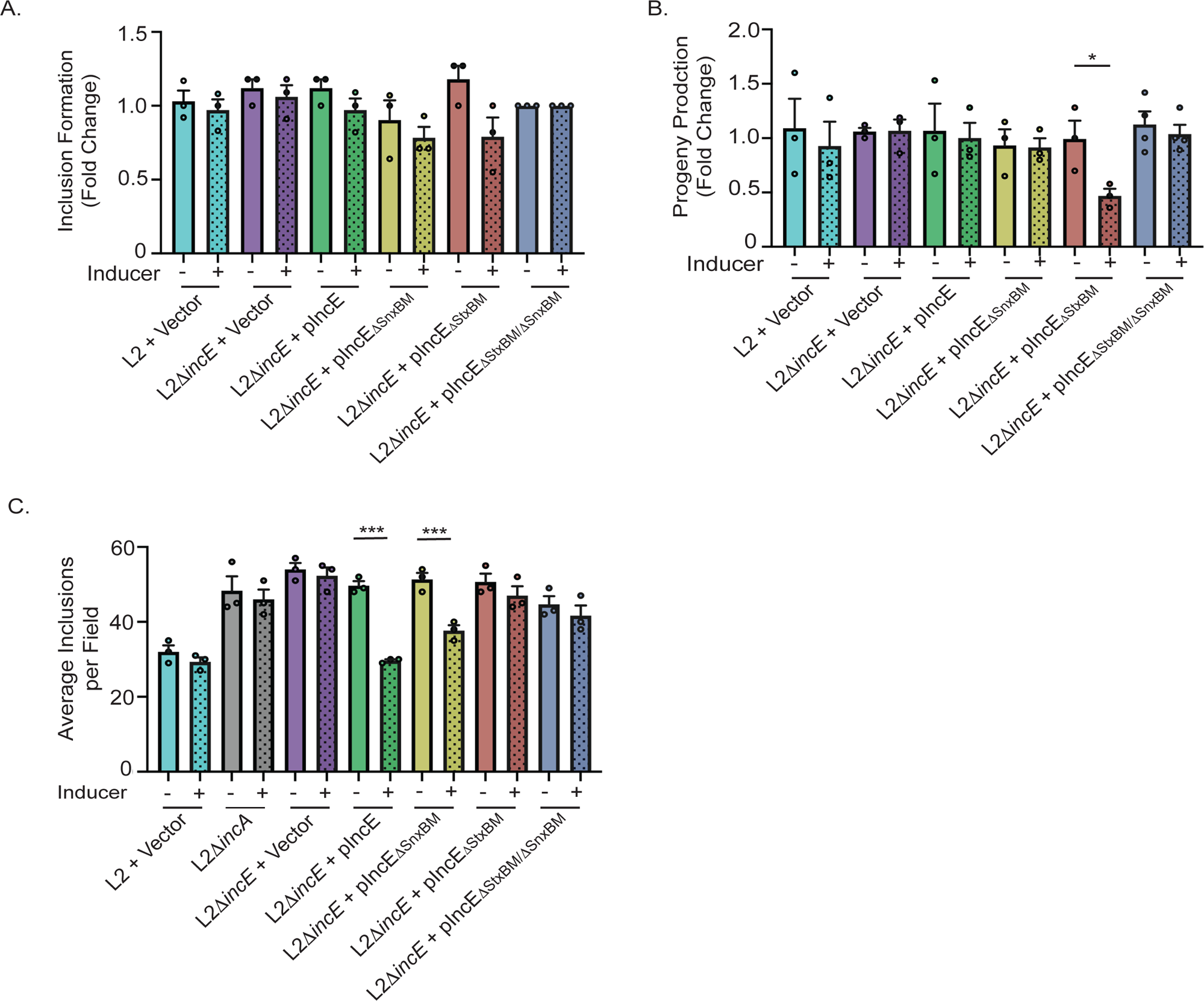
IncE StxBM is required for late steps of infection. Intra-strain comparisons of (A) inclusion formation at 24 hpi and (B) progeny production at 48 hpi. Infections were performed at a low MOI (MOI=1) and normalized to the indicated strain without inducer. (C) Inter-and intra-strain quantification of inclusion fusion at 24 hpi. Infections were performed at a high MOI (MOI=5). Data shown are averages (indicated with circle) ± SEM for at least three biological replicates with at least 99 fields counted for each replicate. * p<0.05, *** p<0.0005, Welch’s ANOVA.

### IncE binding to STX7 and STX12 is required for inclusion fusion

The experiments described above were performed at an MOI ∼ 1, conditions under which most cells are infected with a single EB and therefore start with only a single inclusion. An important aspect of *Ct* development is the ability of multiple inclusions within a single cell to fuse with each other (inclusion fusion), which occurs when cells are infected at a high MOI (van Ooij et al., 1998). Under these conditions, which can be observed in *Ct* human infections (Barnes et al., 1985; Geisler et al., 2001), individual EBs are initially taken up into separate inclusions that subsequently undergo homotypic fusion starting around 12 hpi. Inclusion fusion is thought to be important in the pathogenesis of *Ct* infections *in vivo*. Human genital tract infections with naturally occurring *Ct* variants that fail to undergo fusion due to loss of the inclusion protein IncA exhibit milder pathology (Geisler et al., 2001; Suchland et al., 2000). Likewise, in a mouse genital tract infection model, L2Δ*incA* shows decreased colonization (Suchland et al., 2008).

To assess the role of IncE in inclusion fusion, we infected HeLa cells at a high MOI (MOI=5). Strains lacking functional IncE (L2Δ*incE*+pVector and L2Δ*incE*+pIncE_ΔStxBM/ΔSnxBM_) exhibited decreased inclusion fusion at 24 hpi (Fig. 3C). The delay in fusion was relative, as inclusion fusion approached WT levels at 48 hpi (Fig. S7C). Infection with a mutant that lacks IncA (L2Δ*incA*) (Weber et al., 2016), an Inc that is absolutely required for inclusion fusion (Johnson and Fisher, 2013), failed to undergo inclusion fusion even at 48 hpi (Fig. S7). To define which IncE binding activity was required for efficient inclusion fusion, we compared fusion in cells infected with the *incE* mutant that cannot bind STX7/STX12 (L2Δ*incE*+ pIncE_ΔStxBM_) to cells infected with the *incE* mutant that cannot bind SNX5/SNX6 (L2Δ*incE*+ pIncE_ΔSnBM_). These experiments demonstrated that IncE binding to STX7/12 but not to SNX5/6 is required for efficient inclusion fusion (Fig. 3C, Fig. S7).

### STX7 and STX12 play distinct roles in Ct infection

*Ct* expressing the IncE variant that cannot bind STX7/12 exhibits two different phenotypes—a decrease in progeny production and a delay in inclusion fusion. These two phenotypes are not always linked, as the Δ*incA* mutant, which fails to undergo inclusion fusion, does not exhibit a progeny defect during infection in cultured cells (Weber et al., 2016). Thus, STX7 and STX12 could play distinct roles in *Ct* infection—binding to one STX may be required for efficient inclusion fusion whereas binding to the other STX may be required for progeny production. To test this hypothesis, cells were depleted for STX7 or STX12 by RNAi, infected with L2, and assayed for inclusion formation and production of infectious progeny. Depletion of STX7 (which did not affect STX12 levels; Fig. S8A) resulted in a decrease in progeny production (Fig. 4A) but had no effect on inclusion formation and inclusion fusion (Fig. 4B-D). In contrast, depletion of STX12 (which did not affect STX7 levels; Fig. S8A) resulted in a defect in inclusion fusion at 24 hpi (Figs. 4B-D) but had no statistically significant effect on progeny production (Fig. 4A). Depletion of SNX5/6 had no effect on inclusion fusion (Figs. 4C,D). Together, these results support the hypothesis that binding of IncE to STX7 and STX12 contributes to distinct and separate aspects of *Ct* development. STX7 is required for efficient progeny production while STX12 is required for efficient inclusion fusion. These experiments provide additional support for the observation that a decrease in inclusion fusion does not affect production of infectious progeny.

**Fig. 4.**
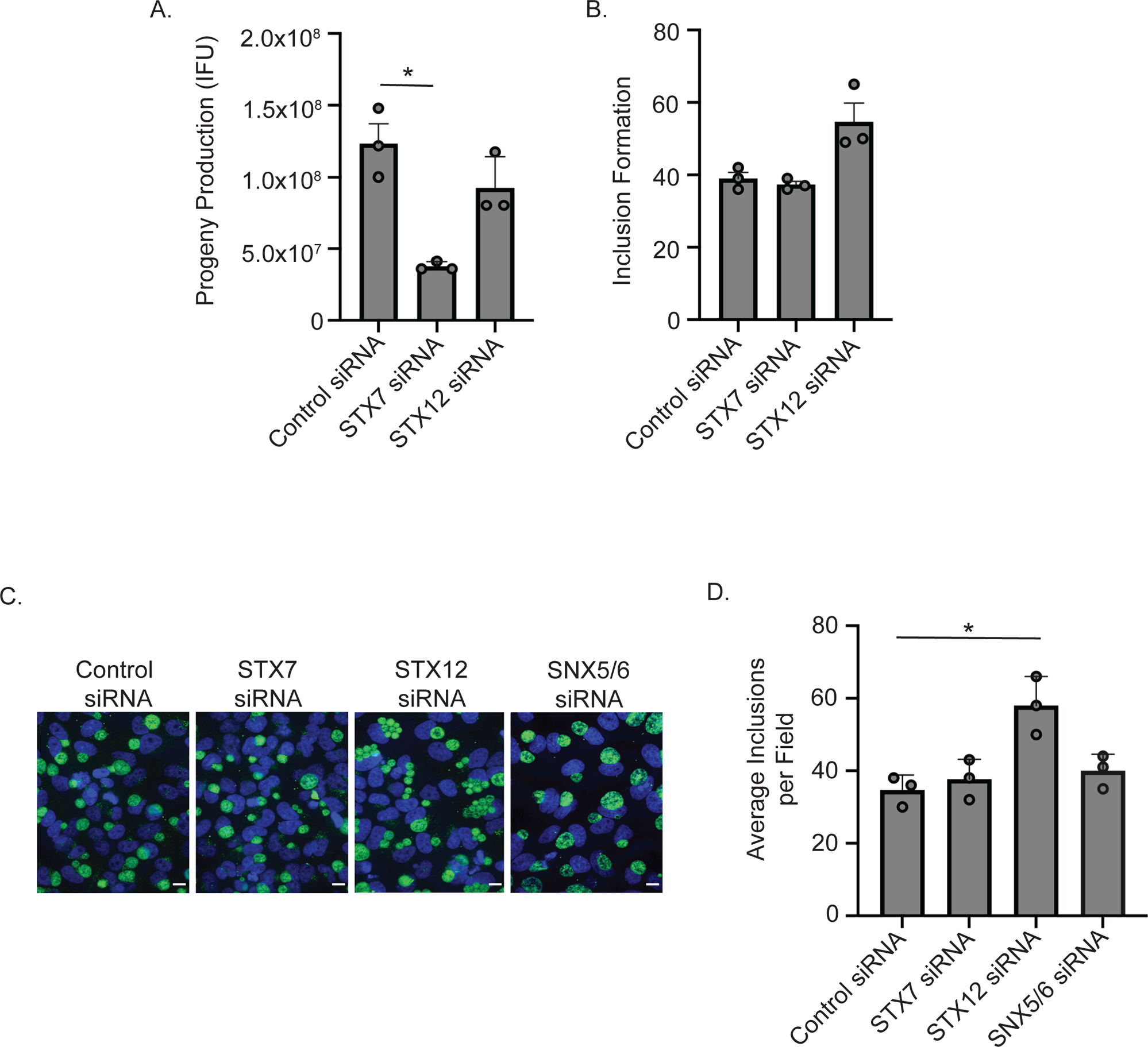
STX7 and STX12 serve distinct functions during infection. Quantification of (A) inclusion number, (B) inclusion size, and (C) production of infectious progeny in control siRNA, STX7 siRNA-, or STX12 siRNA-depleted cells infected (MOI = 0.8) for (A, B) 24 hrs or (C) 72 hrs. Data shown is representative of three biological replicates and is averages ± SEM for three technical replicates with a total of 33 fields counted for each replicate. (D) Representative single slice confocal Immunofluorescence images (scale bar = 10 μm) and (E) quantification of homotypic inclusion fusion in control, STX7-, STX12-, or SNX5/6-depleted cells infected (MOI = 5) for 24 hrs. Multiple, unfused inclusions are present in the STX12 siRNA-treated sample. (E) Data shown are averages ± SEM for three biological replicates with a total of 99 fields counted. The decreased inclusion fusion in the STX12 siRNA-treated cells is reflected by an increase in the average number of inclusions per field and decrease inclusion size. * p<0.05, ** p<0.005, Welch’s ANOVA.

### IncE interacts simultaneously with STX7/12 and SNX5/6 to enhance formation of a hybrid class of vesicles at the inclusion

The remarkable compactness of the IncE C-terminus, with 42 amino acids encoding two non-overlapping motifs that interact with proteins associated with distinct classes of vesicles, raised the possibility that such a reductionist architecture could enable simultaneous and synergistic interactions of IncE with SNX5 or SNX6 and with STX7 or STX12. To explore this notion, we determined whether IncE could co-AP simultaneously SNX5 or SNX6 with STX7 or STX12 in the context of infection. In the absence of infection, endogenous STX7 or STX12 did not co-AP with transfected SNX5_FLAG_ or mCherry-SNX6 (Figs. S9A,B), confirming that these SNARES do not normally interact with SNX5 or SNX6. However, upon infection with L2, endogenous STX7 or STX12 co-AP’d with transfected SNX5_FLAG_ or mCherry-SNX6 (Figs. S9A,B). This interaction was IncE-dependent, as it was abolished upon infection with the *incE* mutant (L2Δ*incE*). We conclude that IncE can interact simultaneously with SNX5 or SNX6 and with STX7 or STX12.

We investigated the potential biological consequence(s) of the ability of IncE to interact simultaneously with STX7 or STX12 and with SNX5 or SNX6 at the inclusion membrane. SNX5 and SNX6 can be recruited to the inclusion membrane (Mirrashidi et al., 2015) and SNX6 is present on vesicles close to the inclusion ((Mirrashidi et al., 2015) see also Fig S5). Therefore, we examined whether STX7 and STX12-containing vesicles are recruited to the inclusion and whether these vesicle fuse with the inclusion. Using live cell imaging of *Ct*-infected cells, we observed GFP-STX7 and GFP-STX12-containing vesicles close to and sometimes fusing with the inclusion membrane (Movies 1 and 2), though these fusion events appeared infrequent.

Given these limited fusion events, we considered the alternative possibility that simultaneous binding of IncE to STX7 or STX12 and to STX5 or STX6 promotes the fusion of SNX5 or SNX6-containing vesicles with STX7 and/or STX12-containing vesicles. Such an event would bring together two normally distinct classes of vesicles in close proximity to potential fuse to form hybrid vesicles. This notion predicts that such hybrid vesicles would be observed proximal to but not distal to the inclusion and that their formation would be IncE-dependent. To test this hypothesis, we examined whether GFP-STX7 and/or mCh-STX12 co-localized with endogenous SNX6 in vesicles that were either proximal or distal to the inclusion. For this quantitation, we determined the location of maximum fluorescence intensity of the 3 proteins in each of 30 vesicles proximal to (within 1 micron) and 30 vesicles distal to (> 5 microns) the inclusion for each experimental group (Fig. 5A). We assessed their overlap the by quantifying distance between their maximal fluorescence intensity.

**Fig. 5.**
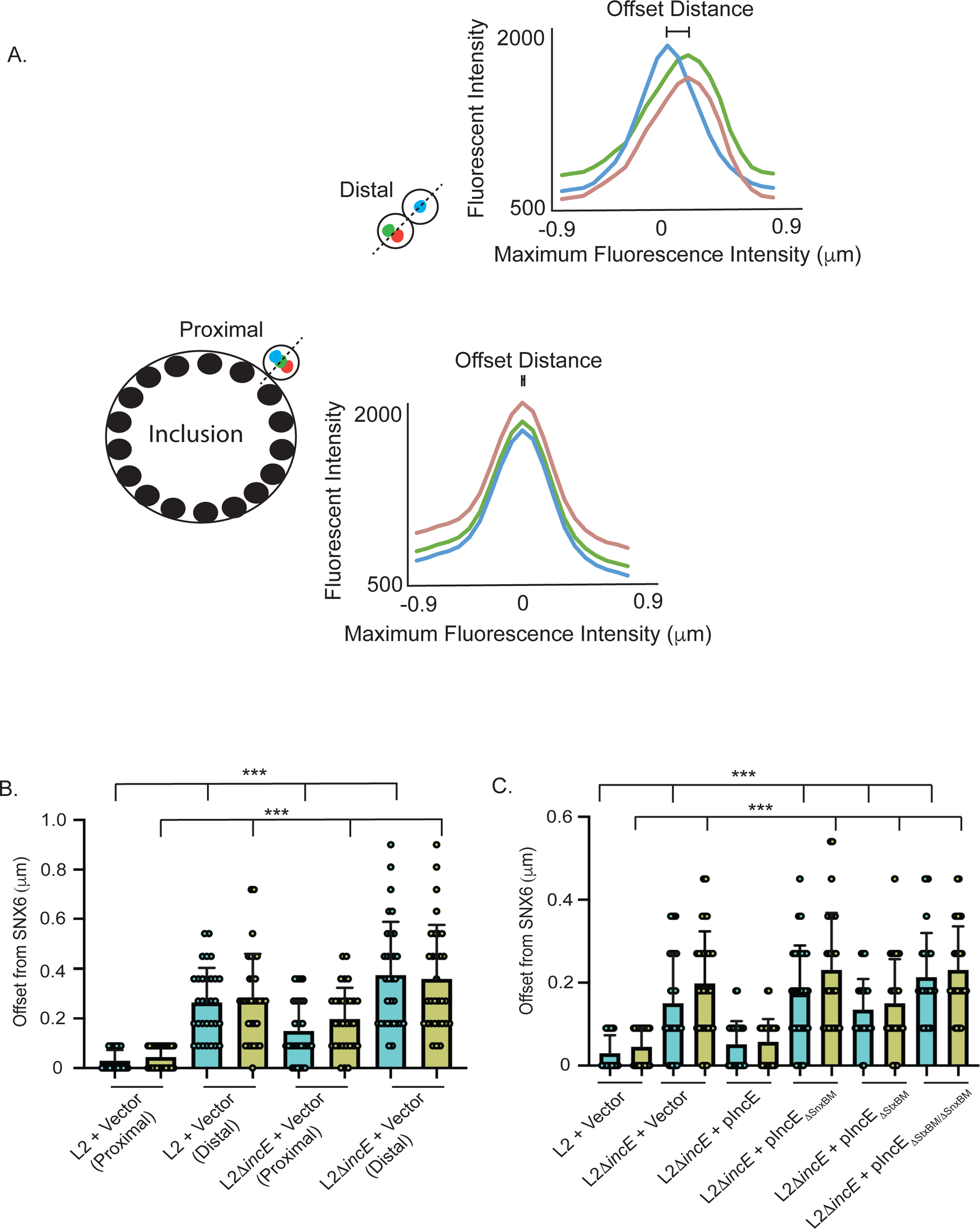
IncE is required for the formation of hybrid SNX6/STX7/STX12-containing vesicles proximal to the inclusion. (A) Schematic depicting quantitation of STX7/STX12/SNX6 overlap in vesicles proximal and distal to inclusion. Fluorescence profiles on vesicles proximal (≤1 micron) and distal (>5 microns) to the inclusion were plotted for GFP-STX7 (green dot), mCh-STX12 (red dot), and endogenous SNX6 (blue dot). The maximum fluorescent intensity offsets of GFP-STX7 and mCh-STX12 from SNX6 were computed. (B) Quantification of STX7/SNX6 (blue) and STX12/SNX6 (yellow) colocalization in vesicles proximal or distal to inclusion in cells infected for 24 hrs with (B) L2+Vector or L2DincE+Vector, or in vesicles proximal to the inclusion in cells infected with (C) L2+Vector, L2D*incE*+Vector, and the indicated L2D*incE*+pIncE variants. All experiments were performed in the presence of inducer. Shown are individual data points as well as the average ± SD for 30 vesicles for each infection condition. *** p<0.0005, Welch’s ANOVA.

In vesicles distal to the inclusion, the maximum fluorescence intensity of GFP-STX7 and of mCh-STX12 was distinct from and did not overlap with endogenous SNX6. The average maximum fluorescence intensity of GFP-STX7 or mCh-STX12 was displaced 0.26 μm and 0.29 μm from the maximum fluorescence intensity of SNX6, respectively (Figs. 5B,C, Table S1). In contrast, the maximum fluorescence intensity of GFP-STX7 and mCh-STX12 often overlapped with each other (Table S1). This result is not entirely unexpected, since both STX7 and STX12 are reported to localize to endosomes, although STX7 is reported to localize to late endosomes (Mullock et al., 2000) and STX12 is reported to localize to early endosomes (Tang et al., 1998). Importantly, in vesicles proximal to the inclusion, the maximum fluorescence intensity of GFP-STX7 and mCh-STX12 overlapped with that of endogenous SNX6 (Figs. 5B,C, Table S1). This finding suggests that formation of hybrid vesicles, which contain STX7, STX12, and SNX6, might be IncE dependent. We tested this prediction explicitly by quantitating the overlap of GFP-STX7, mCherry-STX12, and endogenous SNX6 in vesicles proximal and distal to inclusions in cells infected with L2Δ*incE*. Indeed, the occurrence of these hybrid vesicles proximal to the inclusion was abrogated in the *incE* mutant, and this loss could be rescued by complementation with WT IncE (Figs. 5B,C, Table S1). Similar to infection with L2, there was also no overlap in vesicles distal to the inclusion (Fig. 5B, Table S1). Cells infected with either single or double IncE mutants (*incE*_Δ*StxBM*_, *incE*_Δ*SnxBM*_, *incE*_Δ*StxBM/*Δ*SnxBM*_) displayed no overlap of GFP-STX7 or mCh-STX12 with SNX6 in inclusion-proximal vesicles (Figs. 5B,C, Table S1). While we expect that SNX5 containing vesicles would exhibit similar fusion events with STX7- and STX12-containing vesicles, the absence of an antibody that recognizes endogenous SNX5 precluded testing this prediction directly. We conclude that formation of these hybrid vesicles is IncE-dependent and requires the binding activity of both the StxBM and the SnxBM.

## Discussion

In this work, we demonstrate that IncE is a remarkably compact scaffold that simultaneously interacts with a subset of host proteins associated with distinct vesicle classes to redirect host cell vesicular trafficking to facilitate the *Ct* intracellular life cycle. We and others have previously shown that SNX5 and SNX6 are recruited to the inclusion by binding to a short motif (the SnxBM) present in the C-terminus of IncE. In doing so, the SnxBM of IncE displaces native SNX5/6 cargo, such as the MPR (Elwell et al., 2017; Mirrashidi et al., 2015; Sun et al., 2017), providing a more permissive environment for late steps in the *Ct* intracellular life cycle (Fig. S10). Here, we reveal that IncE encodes a separate non-overlapping 6 amino acid motif (the StxBM) upstream of the SnxBM that mimics the −1 and 0 (ionic) layer of R-SNAREs. The StxBM enables IncE to bind to a subset of Q-SNARES, STX7 and STX12, during infection. Our studies link STX12 with homotypic inclusion fusion and STX7 for inclusion growth and production of infectious progeny. The presence of these two motifs enables IncE to simultaneously bind to STX7 or STX12 and to SNX5 or SNX6, which promotes the formation of hybrid vesicles proximal to the inclusion. This finding suggests that IncE functions in an inclusion-autonomous manner, as would be expected for a membrane-bound effector. Importantly, we demonstrate that the IncE:STX interactions are critical for intracellular growth, progeny production, and homotypic inclusion fusion. We speculate that IncE-dependent tethering and fusion of normally distinct vesicle classes at the inclusion creates a nearby reservoir of accessible nutrients, proteins, and/or lipids that contribute intracellular growth. Future studies will focus on determining the nature of the STX- and SNX-containing vesicles, the identity of the host constituents contributed by these vesicles, and the potential crosstalk between the SNXs and STXs.

We propose that the IncE SnxBM and the StxBM are examples of bacterial effector-encoded short linear motifs (SLiMs). SLiMs are composed of 3-15 amino acids, often embedded in intrinsically disordered or coiled-coil regions. SLiMs adopt a more structured conformation that serves as a binding interface to a structured (globular) partner (Kumar et al., 2022). The human proteome is estimated to encode millions of SLiMs, (Tompa et al., 2014), and SLiMs are also found in viral proteins (Davey et al., 2011; Madhu et al., 2022) and occasionally in bacterial effectors(Samano-Sanchez and Gibson, 2020). The importance of the two SLiMs encoded in IncE is highlighted by the fact that the VLE (found in the StxBM) and VQF (found in the SnxBM) motifs are conserved, albeit with some variation, in several of the IncE homologs found in multiple *Chlamydia* species (Fig S3C). We also note that the arginine in the 0-layer of the R-SNARE is changed to an asparagine (L2), an aspartic acid (Serovars D and B), and a histidine (*C. muridarum* and *C. suis*). How this impacts IncE binding to STX7 or STX12 is unclear; some substitutions for the conserved arginine have been reported to still be functional for SNARE-mediated fusion (Katz and Brennwald, 2000).

The use of SLiMs by Incs to encode a protein-protein binding surface empowers these inclusion-anchored proteins to disrupt, relocate, and/or add functions to host cell complexes. Remarkably, IncE accomplishes all 3 of these tasks: (i) The SnxBM disrupts SNX5/6 binding to the MPR and possibly to other SNX5/6 cargo (Elwell et al., 2017); (ii) Both the SnxBM and the StxBM recruit vesicles to the inclusion; and (iii) By recruiting these 2 distinct classes of vesicles simultaneously, IncE facilitiates the formation of a new hybrid class of vesicles.

We anticipate that the use of SLiMs may be a recurring theme in Inc biology and suggest that SLiMs could account for several unique aspects of Inc biology. Several Incs have been reported to encode their host protein binding interfaces within a short unstructured region or within coiled-coiled sequences in their cytosolically exposed C-termini (Elwell et al., 2017; Sherry et al., 2022; Stanhope et al., 2017). As Incs are rapidly evolving (Lutter et al., 2012), the use of SLiMs would allow rapid diversification of Inc-encoded protein binding domains. Indeed, SLiMs are predicted to evolve *ex nihilo* and to evolve rapidly since they require only a few key amino acids to create a binding surface (Davey et al., 2015; Neduva and Russell, 2005).

Altogether, the C-terminus of IncE is an extraordinarily compact multitasker. It illustrates the vast potential of Incs to modulate and even reprogram the hostile intracellular environment of the host into a mileu that is conducive for the success of this obligate intracellular pathogen. These studies exemplify the power of mining the *Ct*-host interactome for new insights into pathogen virulence strategies as well as for new insights into host cell biology.

## Acknowledgements

We thank Drs.Ted Hackstadt, Dan Rockey, Isabelle Derré, Sourav Bandyopadhyay, and Kenneth Fields for reagents. This work was supported by grants from the NIH (NIAID RO1AI1163526, RO1AI122747, R56AI152526 to JE and NIAID F32AI133902 to KP)

## Author contributions

Conceptualization: KP, KM, CE, JE, JS, EM, CH

Methodology: KP, JS, CT, CH, EM

Investigation: KP, KM, CT

Visualization: KP, CH, CE, JE

Funding acquisition: JE, KP

Supervision: KP, CE, JE

Writing – original draft: KP, CE, JE

Writing – review & editing: KP, CE, JE

## Competing interests

Authors declare that they have no competing interests.

## STAR Methods

### Cell culture and bacterial propagation

HeLa and Vero cells were obtained from American Type Culture Collection (ATCC). HeLa cells were cultured and maintained in Eagle’s Minimum Essential Medium (MEM; UCSF Cell Culture Facility) supplemented with 10% (v/v) fetal bovine serum (FBS) from Gemini at 37°C in 5% CO_2_. HEK293T cells were a generous gift from Nevan Krogan (University of California, San Francisco). HEK293T and Vero cells were cultured and maintained in Dulbecco’s modified Eagle’s Medium (DMEM, UCSF Cell Culture Facility) supplemented with 10% (v/v) FBS at 37°C in 5% CO_2_. Cells were routinely tested for mycoplasma (Molecular Probes, M-7006). *C. trachomatis* L2 (434/Bu; hereafter referred to as L2) was a generous gift from Deborah Dean (University of California, San Francisco). L2 and derivative strains used in these studies are listed in Table S1. *C. trachomatis* was routinely propagated in Vero cell monolayers as previously described (Elwell et al., 2011). HeLa cells were used for all infection studies, and HEK293T cells were used for all ectopic expression experiments. Stellar chemically competent *Escherichia coli* (Takara Bio) were used to propagate constructs for ectopic expression in mammalian cells, while 10-beta *E. coli* (NEB) and dam^−^/dcm^−^ chemically competent *E. coli* (NEB) was used to propagate constructs for transformation into *C. trachomatis*.

### Antibodies and reagents

Primary antibodies were obtained from the following sources: mouse anti-STX7 (Santa Cruz Biotechnology Inc, sc-514157), rabbit anti-STX12 (Atlas Antibodies, HPA055300), rabbit anti-STX2 (Abcam, AB12369-1001), rabbit anti-SNX5 (Proteintech, 17918-1-AP), mouse anti-SNX6 (Santa Cruz Biotechnology Inc, sc-365965), goat anti-SNX6 (Santa Cruz Biotechnology, sc-8679), mouse anti-FLAG (Sigma, F3165), rabbit anti-FLAG (Sigma, F7425), mouse anti-GAPDH (Millipore, MAB374), goat anti-MOMP L2 (Fitzgerald, 20C-CR2104GP), rabbit anti-Strep TagII HRP (Millipore, 71591–3), mouse anti-GFP (Roche, 11814460001), rabbit anti-RFP (Rockland, 600-401-379-RTU). Mouse anti-IncA and rabbit anti-IncG antibodies were kindly provided by Dan Rockey (Oregon State University) and Ted Hackstadt (Rocky Mountain Laboratories), respectively. Rabbit anti-IncE antibody generated against the peptide CKSSPANEPAVNFFKGKNGS (corresponding to amino acids 102–121) was kindly provided by Ted Hackstadt or prepared commercially by Genscript. Secondary antibodies for immunofluorescence were derived from donkey and purchased from Life Technologies or Abcam: anti-goat Alexafluor 647, anti-mouse Alexafluor 647, anti-rabbit Alexafluor 647, anti-mouse Alexafluor 568, anti-rabbit Alexafluor 568, anti-rat Alexafluor 568, anti-goat Alexafluor 488, anti-mouse Alexafluor 488, anti-rabbit Alexafluor 488, anti-rabbit Alexafluor 405, anti-mouse Alexafluor 405. Heparin sodium salt was purchased from Sigma (H3393). Primers (Table S2) were commercially generated by Integrated DNA Technologies or by Elim Biopharm. Dilutions of primary antibodies for immunoblotting and immunofluorescence were performed as follow: anti-Strep (1:1,1000), anti-FLAG (1:3,000 and 1:200), anti-GFP (1:3,000), anti-RFP (1:3,000), anti-STX7 (1:1,000), anti-STX12 (1:1,000), anti-STX2 (1:1000), anti-SNX5 (1:1,000), anti-SNX6 (1:1,1000), anti-GAPDH (1:20,000), anti-MOMP (1:5,000 and 1:500), anti-IncE (1:500 and 1:100), anti-IncG (1:500 and 1:100), and anti-IncA (1:500 and 1:100).

### Plasmid construction

The IncG gene, IncE gene and IncE deletion constructs used for ectopic expression studies were PCR amplified from purified genomic L2 DNA and subcloned into the EcoRI and NotI sites in pcDNA4.0/2xStrepII (Jager et al., 2011). To generate the IncE point mutants used for ectopic expression studies, primers harboring point mutations were used to PCR amplify IncE DNA fragments containing the indicated point mutations. The IncE DNA fragments were ligated together by overlapping PCR and IncE mutants were subcloned into the EcoRI and NotI sites in pcDNA4.0/2xStrepII. STX7 and STX12, and various deletion derivatives were PCR amplified from STX7 and STX12 plasmids (a kind gift of Dr. Sourav Bandyopadhyay, University of California, San Francisco) (Yang et al., 2011) and subcloned into the HindIII and KpnI sites in pEGFP C1 and pmCherry C1 vectors. To generate pSUmC-*incE*-*loxP*-*aadA-gfp*, 3kb DNA fragments from up- and downstream of *incE* were PCR amplified separately from genomic L2 DNA and subcloned into the SalI and SbfI sites, respectively, in pSUmC-*loxP*-*aadA*-*gfp* (a kind gift of Dr. Kenneth Fields, University of Kentucky) (Keb et al., 2018) by Gibson assembly. The *E. coli/Chlamydia* shuttle vector p2TK2-mCherry (a kind gift of Dr. Isabelle Derre, University of Virginia) (Agaisse and Derre, 2013) was modified by removing the *mcherry* gene by InFusion cloning to generate p2TK2. For plasmids that were transformed into *C. trachomatis* L2 strains, IncE variants were PCR amplified from the corresponding IncE-encoding plasmid and cloned into EagI and SbfI sites in the p2TK2. All cloning were verified by forward and reverse sequencing.

### APs

For StrepTactin affinity purifications, 6 x 10^6^ HEK293T cells were seeded in each two to three 10 cm^2^ plates. Cells were transfected using Continuum Transfection Reagent (GeminiBio), following the manufacturer’s instructions. At 24 hours post transfection, cells were scraped in PBS, pelleted, lysed in 1 mL of ice-cold Lysis Buffer (50 mM Tris pH 7.4, 150 mM NaCl, 1 mM EDTA, 0.5% Igepal (Sigma)) at 4°C for 20 minutes while rotating, and centrifuged at 14,000 RPM at 4°C for 20 minutes. Lysates were incubated with 60 mL of Strep-Tactin Sepharose beads (IBA) overnight on a nutator (Clay Adams Brand). Beads were washed five times in 1mL Lysis Buffer containing 0.5% Igepal. Samples were eluted in 60 mL of 2.5mM D-desthiobiotin (IBA) in Lysis Buffer for 30 minutes on rocker at room temperature. Eluates were immunoblotted with anti-STX7, anti-STX12, anti-STX2, anti-SNX5, anti-SNX6, anti-Strep-HRP, and anti-GAPDH antibodies.

For other APs, HeLa cells (3 x 10^5^ cells) were seeded in each well of two 6-well plates. Cells were transfected with indicated construct using Continuum Transfection Reagent (GeminiBio) according to the manufacturer’s instructions. At 6-24 hours post transfection, HeLa cells were infected with the indicated L2 strains (MOI 3) by centrifugation at 1000 RPM for 30 minutes at 4°C. Infected cells were incubated at 37°C in 5% CO_2_ for 30 minutes, and infection media was removed. Fresh media containing 50 ng/mL anhydrous tetracycline (aTC, Takara biotech) was added to infected cells. At 24 hours post infection (hpi), cells were scraped in PBS, pelleted, and lysed in 1 mL of ice-cold Lysis Buffer containing 0.5% Igepal at 4°C for 20 minutes while rotating. Lysates were centrifuged at 14,000 RPM at 4°C for 20 minutes. For FLAG APs, lysates were incubated with 60 mL of anti-FLAG M2 Magnetic Beads (Sigma) overnight on the nutator. Beads were washed five times in 1mL Lysis Buffer containing 0.5% Igepal. Samples were eluted with 150 mg FLAG peptide (Sigma) in Lysis Buffer for 30 minutes on a rocker at room temperature. For mCherry and GFP APs, lysates were incubated with 10 µg rabbit anti-RFP and mouse anti-GFP antibodies, respectively, overnight on the nutator. Lysates were then incubated with Protein G Agarose beads (Pierce) on the nutator at 4°C for 3 hours. Beads were washed five times in 1mL Lysis Buffer. Samples were eluted with 3X LDS Sample Buffer. Eluates were immunoblotted with anti-STX7, anti-STX12, anti-STX2, anti-SNX5, anti-SNX6, anti-Strep-HRP, anti-RFP, anti-GFP, anti-MOMP, and anti-GAPDH antibodies.

### siRNA depletion studies

HeLa cells were transfected with SmartPool siRNA (Dharmacon) for STX7, STX12, SNX5 and SNX6, or siGENOME pool non-targeting #2 using Dharmafect following the manufacturer’s instructions (Dharmacon). At 72 hours post transfection, cells were infected with L2 (MOI= 0.8 or 5) by centrifugation at 1000 RPM for 30 minutes at 4°C followed by incubation at 37°C in 5% CO_2_ for 30 minutes. Infection media was removed, and fresh media was added to infected cells. At 24 hpi and 48 hpi, infected cells were processed for quantitation of inclusion formation or production of infectious progeny, respectively, as described below. siRNA depletion efficiency was determined by immunoblotting lysates with anti-STX7, anti-STX12, anti-SNX5, anti-SNX6, and anti-GAPDH antibodies.

### Fluorescence microscopy

HeLa cells were grown on acid-treated glass coverslips (Warner Instruments) in 24-well plates. For localization of IncE_FLAG_ variants, cells were transfected with GFP-SNX6 using Effectene (Qiagen) following manufacturer’s instructions. For experiments assessing co-localization of STX7, STX12, and SNX6 vesicles, cells were co-transfected with GFP-STX7 and mCherry-STX12 as described above. At 6-24 hours after transfection, cells were infected with indicated L2 strains. Bacteria suspended in MEM supplemented with 10% FBS were centrifuged onto cell monolayers at 1000 RPM for 30 minutes at 4°C. Infected cells were incubated at 37°C in 5% CO2 for 1 hour. Infection media were aspirated, fresh media containing 2 ng/mL aTC were added, and cells were incubated at 37°C in 5% CO2 for 24 hours or 48 hours. Infections for experiments to assay protein localization were performed at MOI 1 in the presence of 2 ng/mL aTC. Experiments assaying homotypic inclusion fusion were performed at MOI 5 in the presence or absence of 2ng aTC. Experiments assaying inclusion formation were performed at MOI 1 in the presence or absence of 2ng/mL aTC. For experiments assaying production of infectious progeny, EBs were harvested from primary infections (as described below) performed at MOI 1 in the presence or absence of 2 ng/mL aTC. The harvested EBs were then used to infect fresh HeLa cells at varying MOIs on coverslips. At 24 hpi, the cells were fixed in 4% PFA in PBS for 15 minutes at room temperature and then permeabilized in 0.1% Triton X-100 in PBS for 15 minutes at room temperature. Cells were blocked in PBS containing 1% BSA for 1 hour and then stained with indicated primary and fluorophore-conjugated secondary antibodies in 1% BSA for 1 hour each. Coverslips were mounted on Vectashield mounting media with or withot DAPI (Vector Laboratories) and imaged on laser scanning disc confocal microscope.

For live cell microscopy, HeLa cells were grown on 24-well glass-bottom plates (MatTek, P24G-1.0-13-F) and co-transfected mCherry-SNX5 as well as either GFP-STX7 or GFP-STX12 using Continuum Transfection Reagent (GeminiBio), following manufacturer’s instructions. At 6 hours after transfection, cells were infected with L2 (MOI=1). At 24 hpi, cells were washed with pre-warmed PBS. Phenol-free DMEM supplemented with 10% FBS and 1mM HEPEs (UCSF Cell Culture Facility) were added and infected cells were imaged on spinning disk confocal microscope as described below.

Images were acquired using Yokogawa CSU-X1 spinning disk confocal mounted on a Nikon Eclipse Ti inverted microscope equipped with an Andora Clara digital camera and CFI APO TIRF 60X oil or PLAN APO 40x objective. Single Z slices were acquired for images used for protein localization, quantifying inclusion formation, inclusion size, production of infectious progeny, and homotypic inclusion fusion. For images used for assessing vesicle co-localization, 0.3 µm-thick Z-stack images were acquired. For live cell imaging, single Z slices were acquired for 60 seconds at 1 frame per second using the CFI APO TIRF 60X oil objective lens. Images were acquired by NIS-Elements software 4.10 (Nikon). For each set of experiments, the exposure time for each filter set for all images was identical. Images were processed with FIJI Software to quantify inclusion numbers and sizes.

### Generation of L2 strains

L2 overexpressing IncE_FLAG_ was generated as previously described (Wang et al., 2011). 10 mg DNA was used to transform 1 x 10^7^ infection forming units (IFUs) of L2 in 200 ul Transformation Buffer (10 mM Tris pH 7.4 in 50 mM CaCl2). Following 30 min incubation at room temperature, the transformation mix was added to 12 mL MEM supplemented with 10% FBS. 2 mL of suspended bacteria was added to each well of 6-well plate containing Vero cells (2 x 10^5^/well). At 12 hpi, 5 mg/mL Ampicillin (Amp; Sigma) and cycloheximide (1 µg/mL, Sigma) was added to select for transformed L2. After 3 initial passages, Amp was increased to 50 mg/mL and L2 transformants were passaged 2-3 times in Vero cells. Clonal populations of transformants were isolated under Amp selection by plaque assay in Vero cells. L2 overexpressing IncG_FLAG_ was a generous gift from Dr. Isabelle Derre (University of Virginia).

Deletion of *incE* in L2 (hereafter referred to L2D*incE*) was generated using floxed-cassette allelic exchange mutagenesis (Keb et al., 2018). L2 was transformed with unmethylated pSUmC-*incE*-*loxP*-*gfp*-*aadA* in Transformation Buffer as described above. At 12 hpi, Spectinomycin (Spec; 500 µg/mL, Sigma), aTC (50 ng/mL; Takara Bio), and cycloheximide (1 µg/mL) were added to select for transformed *Chlamydia*. After 3-4 initial passages, surviving transformants expressing GFP and mCherry were selected by passage in Vero cells in the presence of Spec (500 µg/mL) but without aTC to enrich for plasmid integration by allelic exchange. Clonal populations harboring *incE*::loxP-*aadA-gfp* was plaque-purified in the presence of Spec (500 µg/mL) selection. After 2-3 passages in Vero cells, the plaque-purified transformants were transformed with pSU-CRE, which expresses Cre recombinase, to allow for excision of the *aadA-gfp* cassette on the chromosome. At 12 hpi, Amp (5 mg/mL) and aTC (50 ng/µL) were added to select for transformants. After 3-4 passages, transformants expressing only mCherry (*aadA*-*gfp* cassette excised) were passaged in the absence of Amp and aTC to allow for loss of pSU-CRE to generate L2D*incE*. Loss of incE by Cre-mediated excision at the lox sites was confirmed by PCR screening using a forward primer to the upstream gene (*incD*) and a reverse primer to the downstream gene (*incF*). The presence or absence of the transcripts encoded in the operon (IncD, IncE, IncF, IncG) were confirmed by (i) RT-PCR (primers are listed in Table S1), performed according to the manufacturers protocol (Qiagen) (ii) immunoblot analysis, using antibodies to IncE and IncG, and (iii) immunofluorescence using primary antibodies to IncA, IncE, and IncG. L2D*incE* was transformed with p2TK2 (empty vector) or complemented with the indicated p2TK2-IncE_FLAG_ (IncE_FLAG_) variants. L2Δ*incA*, carrying a *bla* cassette inserted at the *incA* locus, was generated using TargeTron (Johnson and Fisher, 2013).

### Quantitation of inclusion formation

To quantify inclusion formation, HeLa cells infected with the indicated L2 strains for 24 or 48 hours were fixed with 4% PFA, stained with anti-MOMP and fluorescent secondary antibodies, and visualized by spinning disk confocal microscopy. A total of 11 fields per coverslip x 3 (technical replicates) were acquired for a total of 33 fields per condition. Inclusions were quantified using the macro script or Cell Counter on the FIJI software. Data are mean ± SEM of at least 2 independent biological replicates. To quantify production of infectious progeny, infected HeLa cells were osmotically lysed in ddH2O at 48 hpi. 2- to 5-fold serial dilutions of harvested bacteria were used to infect fresh HeLa monolayers. After removal of infection media, fresh media containing 1mg/mL Heparin (Sigma) was added to cells. At 24 hpi, inclusion formation was enumerated as described above.

### Quantitation of protein co-localization

Co-localization of GFP-STX7, mCherry-STX12, and endogenous SNX6 vesicles during *C. trachomatis* infection was quantified by fluorescence microscopy. HeLa cells co-transfected with GFP-STX7 and mCherry-STX12 were infected with indicated L2 strain, fixed, and permeabilized. Cells were incubated with anti-SNX6, anti-IncA, anti-STX12, and fluorescent secondary antibodies. Images were acquired using a 60X objective lens. A total of 10-15 fields per coverslip x 3 technical replicates were acquired. The fluorescence intensity profiles for GFP-STX7, mCherry-STX12, and endogenous SNX6 were generated using the FIJI software. The maximum fluorescent intensity (F_max_) for SNX6 was arbitrarily set to position zero. The maximum fluorescent intensity offsets for STX7 and STX12 from SNX6 were computed and converted to microns (1 arbitrary unit = 0.09 μm) using the following equation:

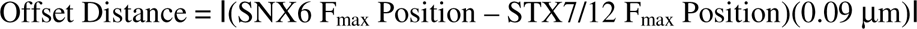

STX7 or STX12 were considered to co-localize with SNX6 when the F_max_ offset distance was ≤0.09µm. STX7 or STX12 were score as not co-localizing with SNX6 is when the F_max_ offset distance was >0.09µm. Data are mean ± SD of 30 vesicles.

### Bioinformatics

The full length IncE protein sequence from *C. trachomatis* L2 (434/Bu) were analyzed using to HMMTOP(Tusnady and Simon, 1998), TMpred(Hofmann and Stoffel, 1993), SOSUI(Hirokawa et al., 1998), and TMHMM(Hofmann and Stoffel, 1993) using standard parameters to predict the two transmembrane on IncE. Multiple sequence alignment of IncE homologs, Q-SNAREs, and R-SNAREs were performed using Clustal Omega (Madeira et al., 2022).

### Statistical analysis

For each experiment, at least 2 or more independent biological/technical replicates were performed, and the results are plotted individually or combined and represented as mean ± SD/SEM, as described in the figure legends. All statistical analyses were performed using GraphPad Prism 9.0. Assays were analyzed using a one-way ANOVA with a two-tailed Welch’s t-test or unpaired Student t-test.

Further information and requests for resources and reagents should be directed to and will be fulfilled by the Lead Contact, Joanne Engel

All unique/stable reagents generated in this study are available from the Lead Contact upon request, but we may require reasonable compensation by the requestor for processing and shipping charges.

## Supplemental Information

Document S1. Figures S1-S10 and Table S1-S2

FIJI Macro Script (for quantitation of inclusion formation and production of infectious progeny)

**Fig. S1. IncE binds specifically to STX7 and STX12.** Co-APs of lysates from (A) HEK293T cells transfected with IncG_Strep_ or IncE_Strep_ or (B) HeLa cells infected with L2+pIncG_FLAG_ or L2+pIncE_FLAG_ in the presence of inducer. STX2 serves as a STX binding control. GAPDH serves as a loading control. MOMP, a *Ct* protein, serves as an indicator of the efficiency of infection.

**Fig. S2. Alignment of the SNARE domains of the Q-SNARE syntaxins.** The heptad repeats (layers) of the SNARE domains are numbered −7 to +7 with the conserved glutamine (Q) in the 0 (ionic) layer highlighted in the red box. Shown on the right is the percent homology of each SNARE domain to the STX7 SNARE domain.

**Fig. S3. Delineation of the IncE StxBM.** (A) Schematic of IncE binding sites for STX7/12 and SNX5/6. The StxBM is colored blue. The V114 and F116 residues critical for IncE binding to SNX5/6 are colored red. (B) Bioinformatic predictions of the IncE transmembrane (TM) regions using HMMTOP, TMpred, SOSUI, and TMHMM. (C) Alignment of the human (Hs) R-SNARE VAMP3 with *Ct* serovar L2, *Ct* serovar D, *Ct* serovar B, *Ct serovar* E, *C. suis*, and *C. muridarum* IncE homologs. The predicted StxBM for each IncE homolog and their alignments with the −1 and 0 layer (VLERDQ of HsVAMP3 and VLERHG of MmVAMP5) are indicated with a dashed line. The IncE TM2 is also depicted. (D) Alignment of IncE, IncE_DStxBM_, and IncE_DSnxBM_ variants. The mutated residues are underlined. (E) Representative immunoblot of lysates and eluates from co-APs of IncE_Strep_ variants. GAPDH serves as a loading control. IncE_ΔSnxBM_ migrates slower than IncE or IncE_ΔStxBM_.

**Fig. S4. Construction of L2**Δ***incE* deletion strains.** (A) Schematic of the *incDEFG* operon for L2, L2D*incE::aadA-gfp*, and L2D*incE*. The regions between *incD*-*incF* targeted for PCR amplification are underlined and the predicted sizes of the PCR product are shown. (B) PCR products of the *incDEF* loci in the indicated strains. (C) RT-PCR confirming mRNA transcripts from the *incDEFG* locus in the indicated strains. IncE transcription is undetectable in the *incE* mutants. IncF and IncG transcripts are decreased in the L2D*incE::aadA-gfp* strain due to polar effect of the inserted *aadA*-*gfp* cassette. (D) Immunoblot of IncE and IncG expression in the indicated strains. IncE protein was undetectable in either mutant. IncG protein is similar to WT levels in the L2Δ*incE* but absent in L2D*incE*::*aadA*-*gfp*. MOMP serves as an internal control for number of bacteria in each sample. (E) Localization of IncE and IncG during infection with the indicated strains. Infected cells were fixed, permeabilized, and stained with anti-IncE, anti-IncG, and anti-IncA (inclusion membrane marker) at 24 hpi. Scale bar, 5 μm. (F) Immunoblot of IncA expression. L2Δ*incE* exhibits wild type levels of IncA. GAPDH serves as a loading control. MOMP serves as an internal control for number of bacteria in each sample.

**Fig. S5. IncE_FLAG_ variants localize to the inclusion membrane during infection.** Confocal immunofluorescence microscopy of HeLa cells transiently transfected with GFP-SNX6 (green) followed by infection with the indicated L2 strains for 24 hrs. Cells were fixed, permeabilized, and stained with anti-FLAG (red), anti-IncA (magenta, inclusion membrane marker), and DAPI (blue). Transfected GFP-SNX6 is found on the inclusion membrane as well as on vesicles on or near the inclusion membrane in all strains except those infected with L2*DincE+*Vector or with the L2 expressing the IncE variants that cannot bind SNX5/6 (pIncE_ΔSnxBM_ and pIncE_ΔStxBM/ΔSnxBM_). N, nucleus. I, inclusion. Scale bar, 5 μm.

**Fig. S6. Loss of IncE has no effect on *Ct* intracellular growth.** (A) Schematic of the *Ct* life cycle. The infectious elementary body (EB; black dot) enters the host cell through receptor-mediated endocytosis and is enclosed in a membrane-bound compartment (inclusion, I) that dissociates from the canonical endolysosomal pathway. The EB differentiates into the non-infectious reticulate body (RB; blue circle) and the inclusion migrates along microtubules (not shown) to the juxtanuclear region. Starting at ∼ 24 hpi, RBs re-differentiate to EBs and exit the host cell at ∼48-72 hpi. N, nucleus. Inclusion formation (number and size of inclusion) is a robust metric of early stage of *Ct* development. Production of infectious progeny additionally assesses late stages of infection and completion of the Ct developmental cycle. (B-D) Quantification of (B) inclusion formation, (C) inclusion size, and (D) production of infectious progeny in cells infected with L2+Vector or L2D*incE*+Vector. Data shown are averages ± SEM for at three biological replicates (indicated with black circles) with at least 99 fields counted per replicate. Unpaired Student t-test.

**Fig. S7. Inclusion fusion of *incE* mutant and *incE* variants recover to wildtype levels at 48 hpi.** Quantification of homotypic inclusion fusion (average inclusion number per field) in HeLa cells infected with L2+Vector, L2D*incE*+Vector, or L2D*incE*+pIncE variants at 48 hpi (MOI 5) in the presence or absence of inducer (aTC). Data shown is the average ± SEM for two biological replicates (individual data points are shown as small circles) with 66 fields counted per replicate. Comparison of each strain in the presence or absence of inducer provides intra-strain comparisons. Comparison between different strains providers interstrain comparisons. Inclusion fusion in strains lacking IncE or lacking the StxBM is similar to WT L2 at 48 hpi, whereas inclusion fusion at 48 hpi is still defective in L2Δ*incA*.

**Fig. S8. Efficacy of RNAi depletion**. STX7, STX12, and SNX5/6 protein levels were assessed by immunoblot analysis following 72 hr RNAi depletion for (A) low MOI (MOI=1) and (B) high MOI (MOI=5). GAPDH serves as a loading control.

**Fig. S9. IncE binds simultaneously to SNX5 or SNX6 and to STX7 or STX12**. Immunoblots of (A) SNX5_FLAG_ and (B) mCh-SNX6 APs. HeLa cells were transiently transfected with (A) SNX5_FLAG_ or (B) mCh-SNX6. Cells were left uninfected (UI) or infected with L2+Vector or L2D*incE*+Vector. At 24 hpi, lysates were AP’d with anti-FLAG magnetic beads (A) or anti-RFP-Protein G agarose beads (B). Eluates were immunoblotted with the indicated antibodies. GAPDH serves as a loading control and MOMP levels correlate with *Ct* infection efficiency.

**Fig. S10. Model of IncE:STX7/STX12 and IncE:SNX5/SNX6 interactions.** IncE encodes two SLiMs, the StxBM and SnxBM, to bind to STX7/12 and SNX5/6, respectively. The IncE SnxBM binds to the SNX5/6 cargo binding domain to reroute SNX6^+^ (grey half-moon; and presumably SNX5^+^ (grey half-moon)) vesicles to the inclusion. By binding to SNX5/6, IncE displaces SNX5/6 cargo proteins and disrupts ESCPE-1-mediated restriction of *Ct* intracellular development. IncE StxBM binding to STX7 (tan swiggle line) and STX12 (tan swiggle line) reroutes STX7- and STX12-containing vesicles to the inclusion. The STX- and SNX-containing vesicles fuse in an IncE-dependent manner. STX7 contributes to the production of infectious progeny, while STX12 functions to promote efficient inclusion fusion.

**Video S1.** Cells were co-transfected with GFP-STX7 and mCherry-SNX5, infected with L2 for 24 hrs, and then imaged every second for 60 seconds by live cell confocal microscopy. Fluorescent vesicles are occasionally observed on or fusing with the inclusion membrane. mCherry-SNX5 serves as marker for the inclusion membrane. Scale bar, 5 μm.

**Video S2.** Cells were co-transfected with GFP-STX12 and mCherry-SNX5, infected with L2 for 24 hrs, and then imaged every second for 60 seconds by live cell confocal microscopy. mCherry-SNX5 serves as marker for the inclusion membrane. Scale bar, 5 μm.

**Table S1. Quantitation of STX7, STX12, and SNX6 co-localization.**

**Table S2. Primers and strains.**

## Notes

### Competing Interest Statement

The authors have declared no competing interest.

