## Supplementary material for "The *Chlamydia* effector IncE employs two short linear motifs to reprogram host vesicle trafficking": Suppl figures 1-10

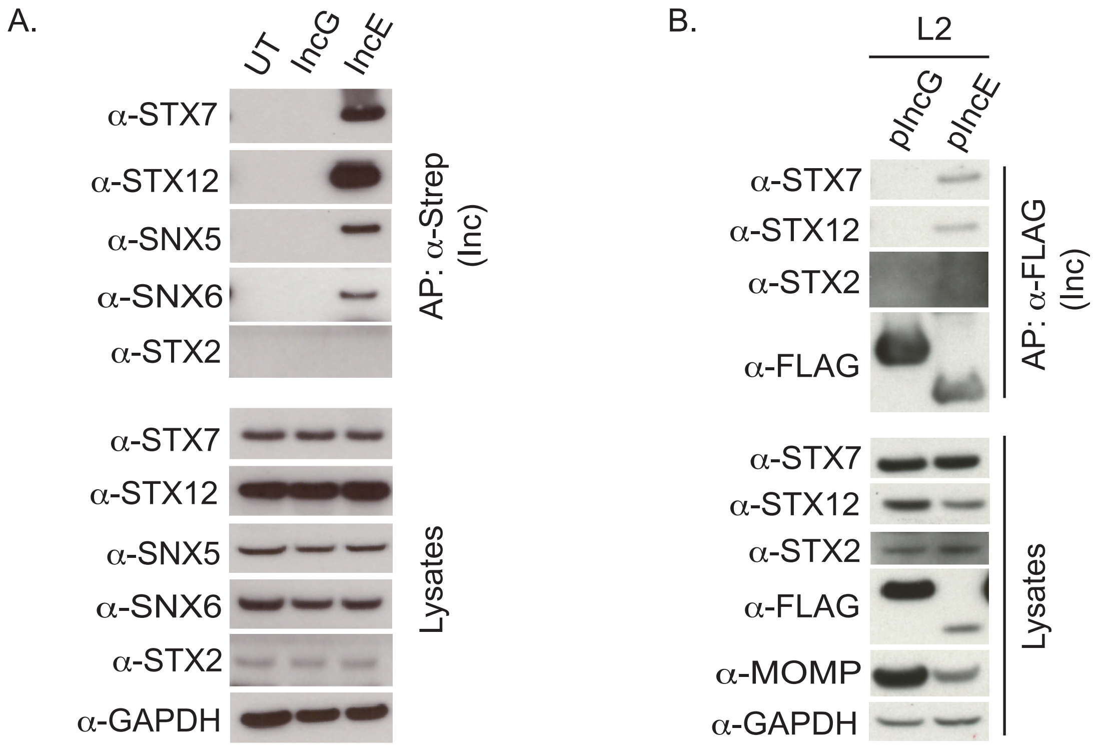


**Fig. S1. IncE binds specifically to STX7 and STX12.**

A.

| | -7 -6 -5 -4 -3 -2 -1 0 +1 +2 +3 +4 +5 +6 +7 | % Homology |
|---|---|---|
| STX7 | IRQLEADIMDINEIFKDLGMMIHEQGDVIDSIEANVENAEVHVQQANQQL | |
| STX12 | IRQLEADILDVNQIFKDLAMMIHDQGDLIDSIEANVESSEVHVERATEQL | 76% |
| STX2 | IMKLETSIRELHEMFMDMAMFVETQGEMINNIERNVMNATDYVEHAKEET | 38% |
| STX3 | IVRLESSIKELHDMFMDIAMLVENQGEMLDNIELNVMHTVDHVEKARDET | 34% |
| STX4 | IQQLERSIRELHDIFTFLATEVEMQGEMINRIEKNILSSADYVERGQEHV | 32% |
| STX5 | MQNIESTIVELGSIFQQLAHMVKEQEETIQRIDENVLGAQLDVEAAHSEI | 30% |
| STX6 | LELVSGSIGVLKNMSQRIGGELEEQAVMLEDFSHELESTQSRLDNVMKKL | 14% |
| STX10 | LEMVSGSIQVLKHMSGRVGEELDEQGIMLDAFAQEMDHTQSRMDGVLRKL | 16% |
| STX8 | LDALSSIISRQKQMGQEIGNELDEQNEIIDDLANLVENTDEKLRNETRRV | 20% |
| STX17 | WETLEADLIELSQLVTDFSLLVNSQQEKIDSIADHVNSAAVNVEEGTKNL | 30% |
| STX18 | VRQIEGRVVEISRLQEIFTEKVLQQEAEIDSIHQLVVGATENIKEGNEDI | 20% |

**Fig. S2. Alignment of the SNARE domains of the Q-SNARE syntaxins.**

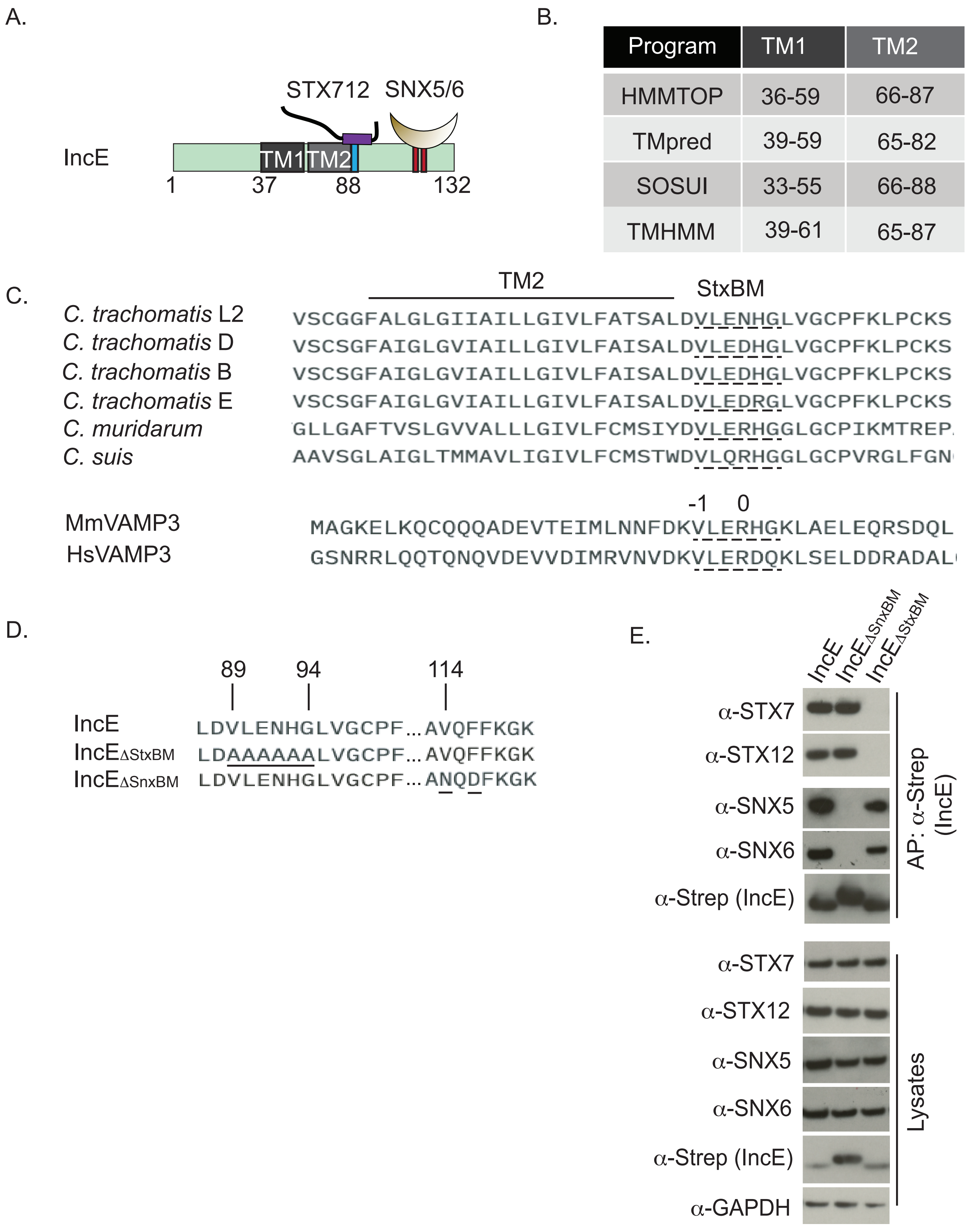


**Fig. S3. Delineation of the IncE StxBM.**

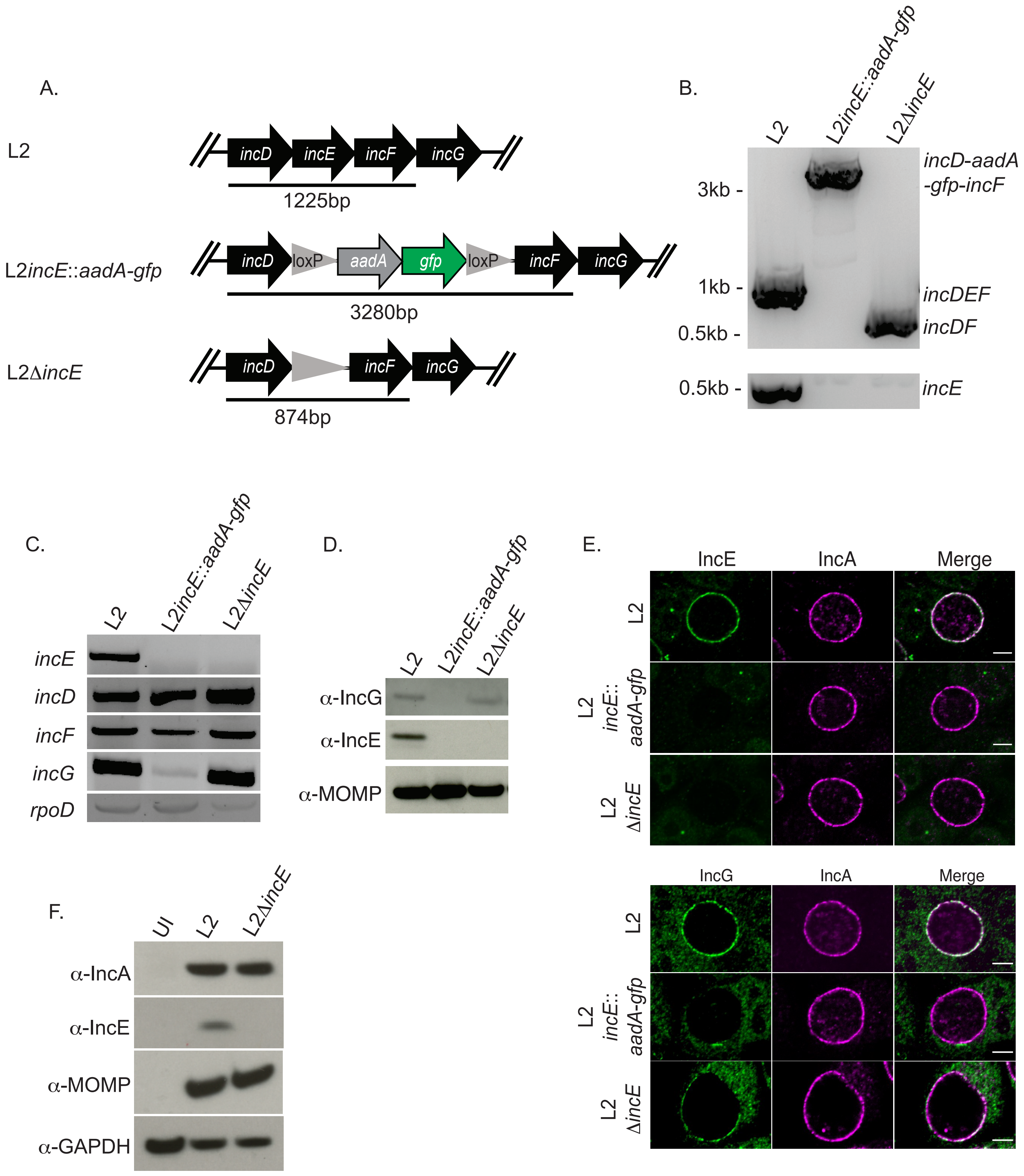


**Fig. S4. Construction of L2Δ*incE* deletion strains.**

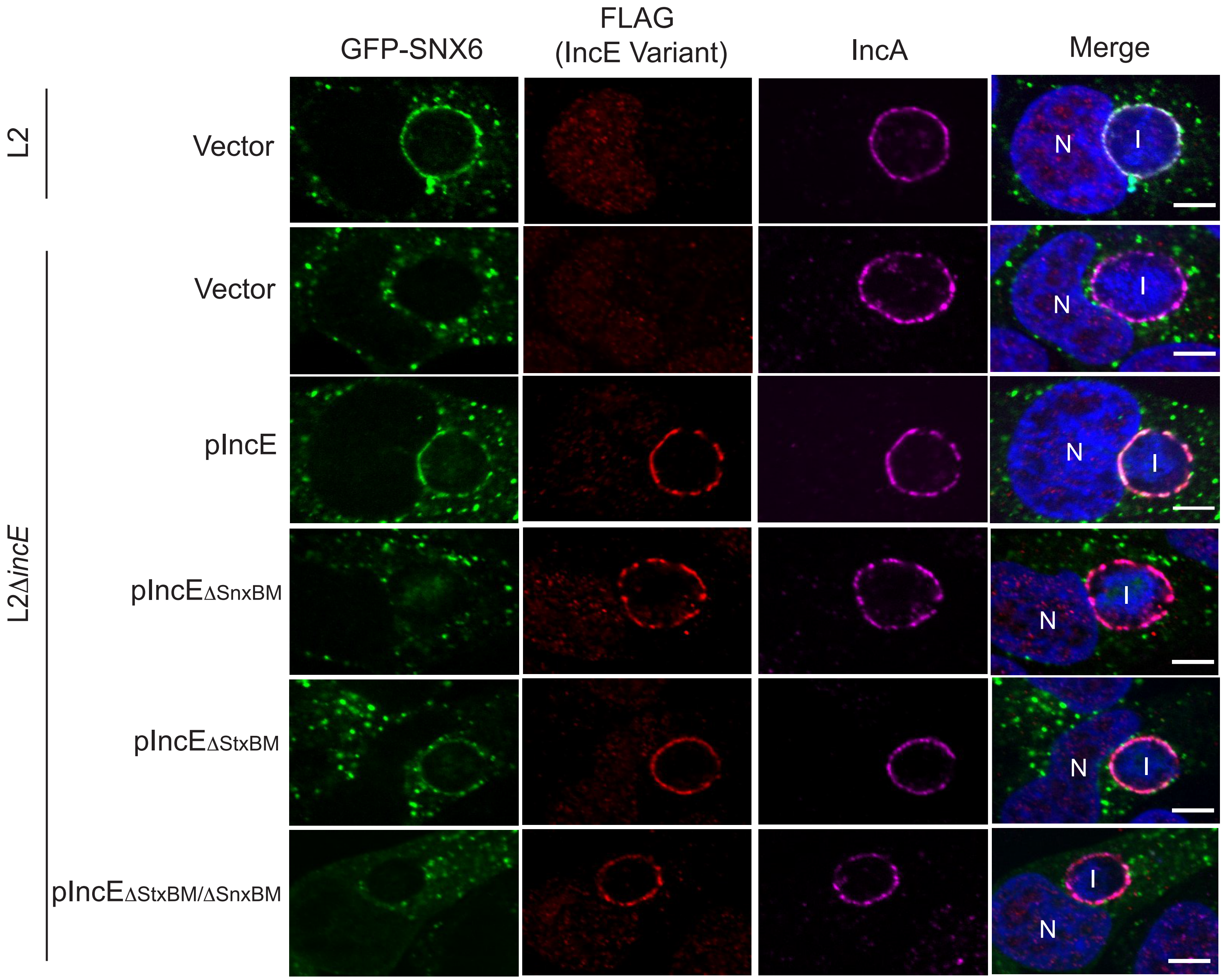


**Fig. S5. IncE$_{FLAG}$ variants localize to the inclusion membrane during infection.**

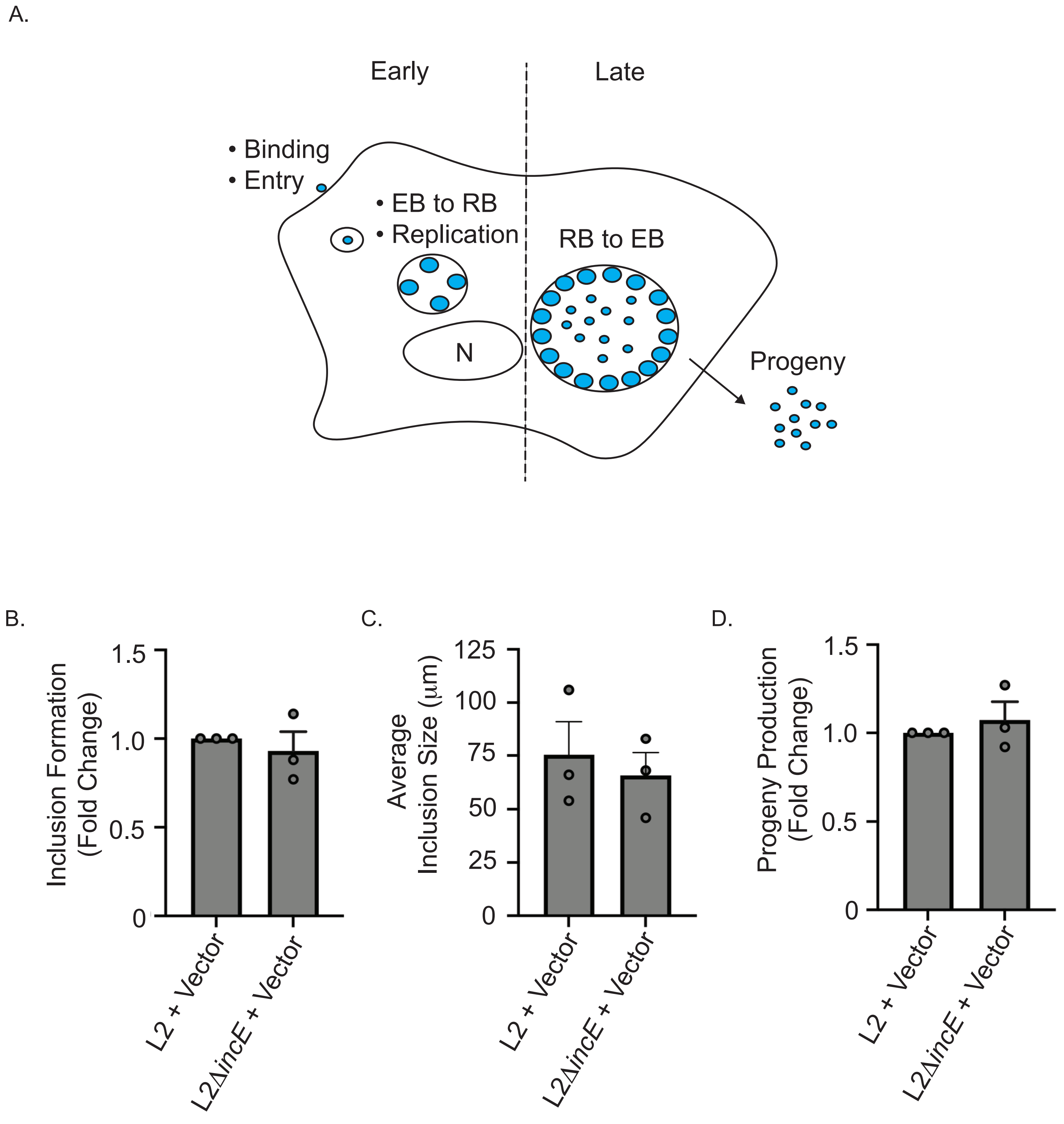


**Fig. S6. Loss of IncE has no effect on Ct intracellular growth.**

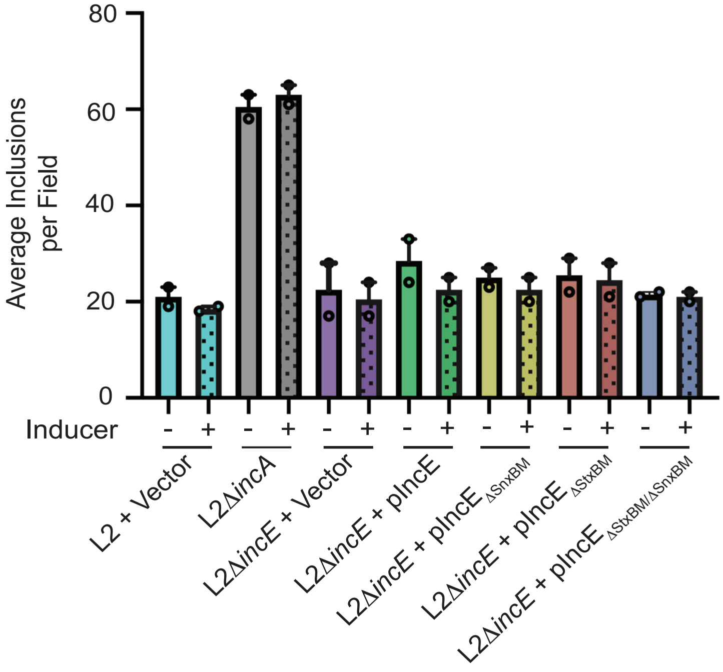


**Fig. S7. Inclusion fusion of *incE* mutant and *incE* variants recover to wildtype levels at 48 hpi.**

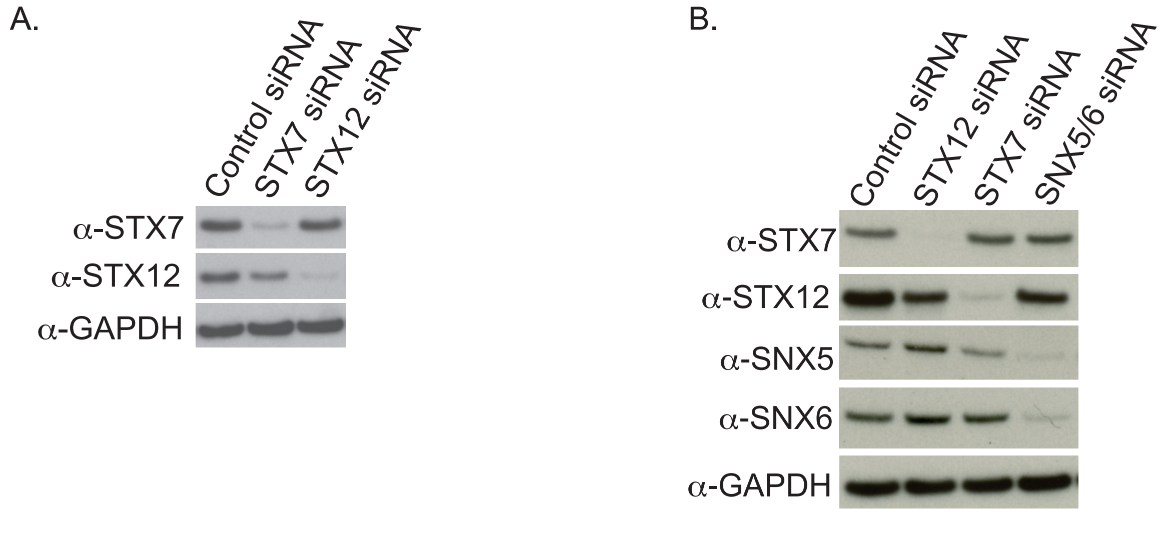


**Fig. S8. Efficacy of RNAi depletion.**

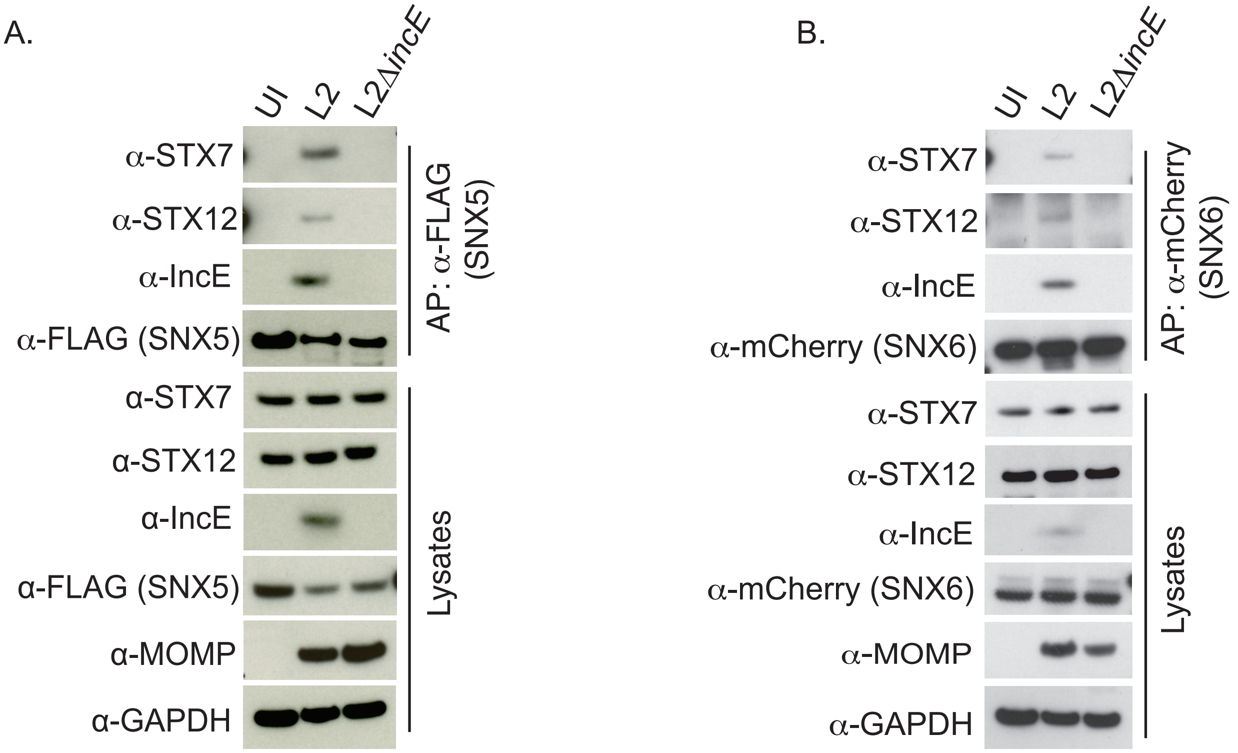


**Fig. S9. IncE binds simultaneously to SNX5 or SNX6 and to STX7 or STX12.**

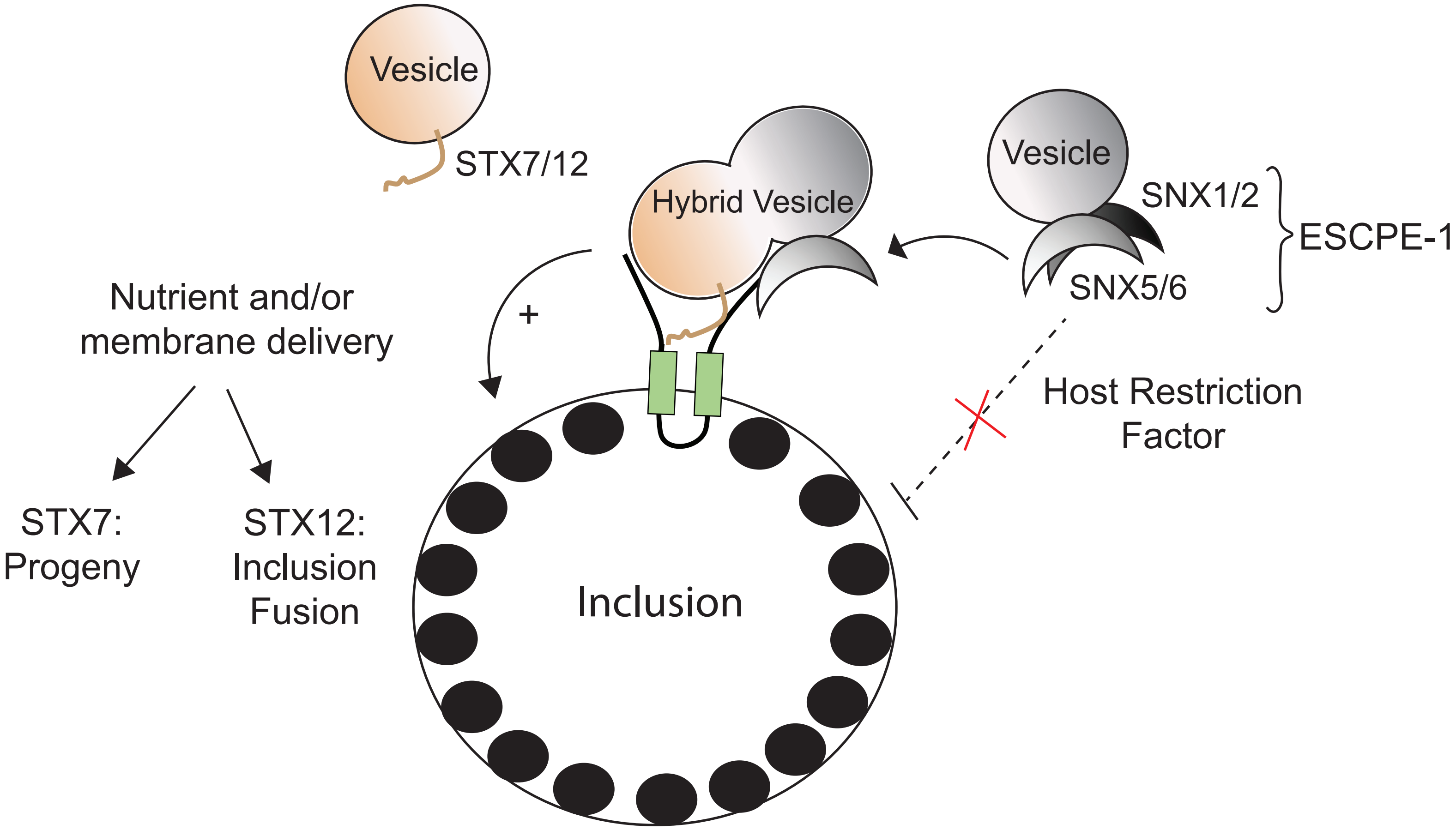


**Fig. S10. Model of IncE:STX7/STX12 and IncE:SNX5/SNX6 interactions.**
